# Endothelial FABP4 constitutes the majority of basal circulating hormone levels and regulates lipolysis-driven insulin secretion

**DOI:** 10.1101/2022.10.13.511807

**Authors:** Karen E. Inouye, Kacey J. Prentice, Alexandra Lee, Carla Dominguez-Gonzalez, Mu Xian Chen, Grace Yankun Lee, Gökhan S. Hotamışlıgil

## Abstract

Fatty acid binding protein 4 (FABP4) is a lipid chaperone secreted from adipocytes upon stimulation of lipolysis. Circulating FABP4 levels strongly correlate with body mass index and obesity-related pathologies in experimental models and humans. While adipocytes have been presumed to be the major source of hormonal FABP4, this question has not been addressed definitively *in vivo*. We generated mice with FABP4 deletion in cells known to express the gene; adipocytes (Adipo-KO), endothelial cells (Endo-KO), myeloid cells (Myeloid-KO), and the whole body (Total-KO) to examine the contribution of these cell types to basal and stimulated plasma FABP4 levels. Unexpectedly, baseline plasma FABP4 was only reduced by ∼25% in Adipo-KO mice, whereas Endo-KO mice showed ∼75% decreases compared to wildtype controls. In contrast, Adipo-KO mice exhibited ∼62% reduction in FABP4 responses to lipolysis, while there was minimal reduction in Endo-KO mice, indicating that adipocytes are the main FABP4 source in lipolysis. We did not detect any myeloid cell contribution to circulating FABP4. Surprisingly, despite the nearly intact FABP4 responses, Endo-KO mice showed blunted lipolysis-induced insulin secretion, identical to Total-KO mice. We conclude that the endothelium is the major source of baseline hormonal FABP4 and is required for the insulin response to lipolysis.

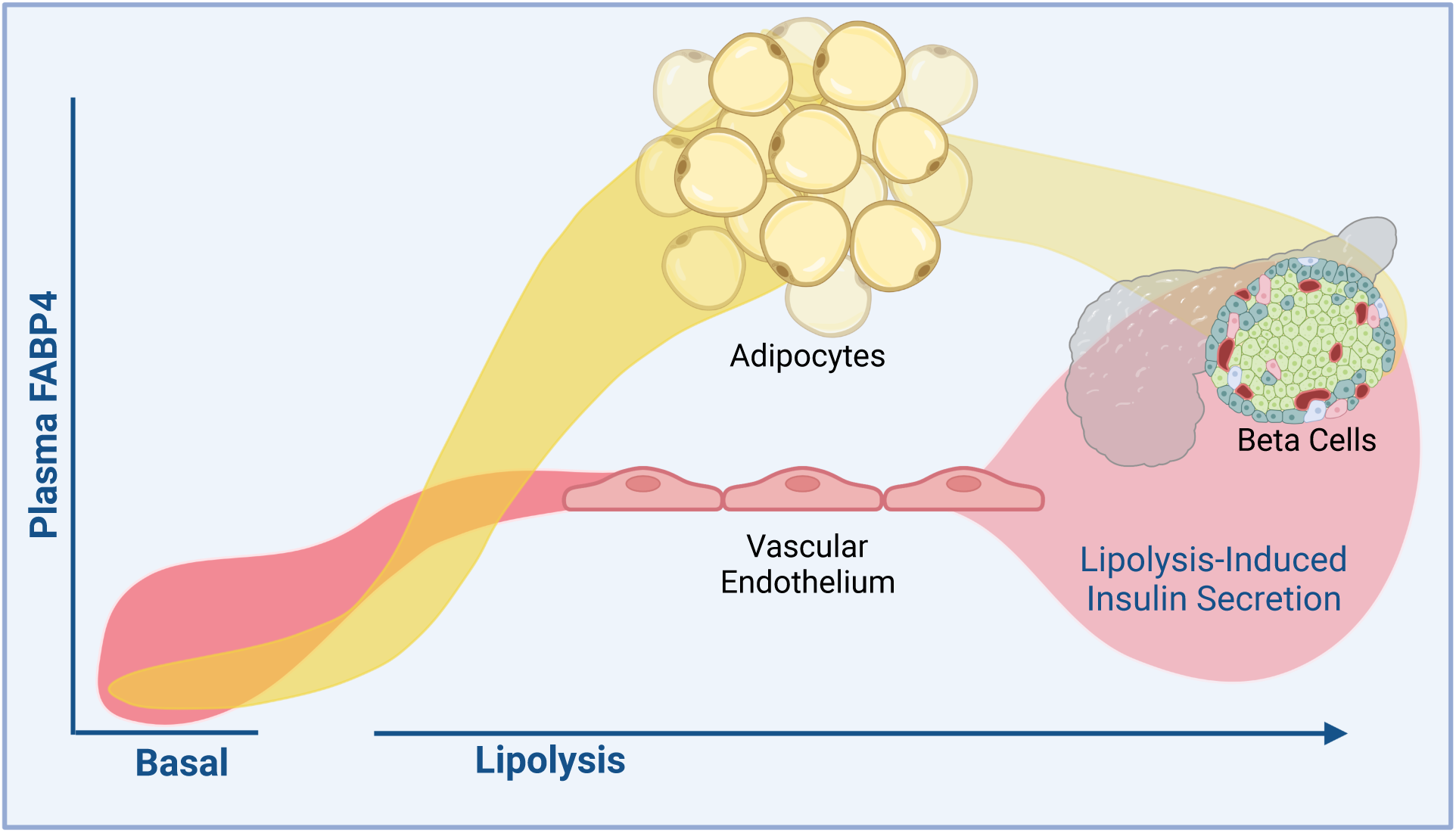

## Introduction

Fatty acid binding protein-4 (FABP4, also known as aP2) is a 15 kDa protein belonging to a highly conserved family of lipid binding proteins, each with specific tissue distributions (1). FABP4 was first identified as the adipocyte-specific isoform, regulated during differentiation, and is one of the most abundant proteins in mature adipocytes, comprising up to 6% of total adipocyte cytosolic protein content (2). Total genetic deletion of FABP4 in mice confers protection against obesity-associated insulin resistance, glucose intolerance, inflammation, fatty liver disease, atherosclerosis, and certain cancers (3–7). Importantly, multiple independent studies in humans carrying a low-expression variant of FABP4 demonstrated lower incidence of type 2 diabetes, decreased circulating triglycerides and cholesterol, and reduced incidence of cardiometabolic disease (8, 9).

FABP4 has been generally considered as a cytosolic protein where it exerts its actions as an intracellular protein. Similar to several other FABP isoforms, it has been detected in serum and this has been ascribed to cell death and leakage from the degenerating source cells (10–12). We have however demonstrated that FABP4 is, in fact, secreted from adipocytes in a regulated manner, through a non-classical pathway, induced by lipolysis (13, 14). At the moment, it is the only known adipocyte hormonal signal accompanying the breakdown of energy stores during lipolysis. Subsequently, it was shown that a major proportion of FABP4 secretion from adipocytes occurs through a lysosomal pathway, a mechanism extremely rare for mammalian protein secretion (15). Circulating FABP4 levels correlate positively with BMI and indicators of metabolic syndrome in humans, and are increased in mice with dietary or genetic obesity (12, 13, 16, 17), raising the possibility that excessive adipose tissue release of FABP4 may in fact be the strong contributor to metabolic dysregulation seen in mouse models as well as in humans (18). Taken together, these observations raise interest in antibody-based targeting of FABP4 hormone for therapeutic purposes, and proof of principle studies in experimental models demonstrated the efficacy and feasibility of this approach (19, 20).

FABP4 secretion by cultured adipocytes and *in vivo* is inducible, stimulated by prolonged fasting and lipolytic signals (13, 14). Induction of lipolysis in mice with the β-adrenergic agonists isoproterenol or CL-316,243 potently stimulates FABP4 secretion (13). It has been shown long ago that acute β-adrenergic stimulation of lipolysis *in vivo* is associated with a rapid, strong, and transient induction of insulin secretion (21, 22). The mechanisms and functional consequences of this rather unusual link have remained enigmatic. Interestingly, studies in FABP4-deficient animals demonstrated that the presence of FABP4 is an important requirement for this lipolysis-driven insulin secretion, as insulin responses to lipolysis are blunted by at least 60% in FABP4-KO mice, whereas they remain intact in response to other signals (23). Most critical to note is that glucose-stimulated insulin secretion from islets of FABP4-deficient animals is enhanced, and that these mice are strongly protected against both type 1 and type 2 diabetes (3, 5, 24). Since FABP4-KO mice are also protected against obesity-associated hyperinsulinemia, the decrease in lipolysis-driven insulin secretion has been considered as one potential contributing factor to their reduced hyperinsulinemia and improved metabolic health (3, 5, 23).

While FABP4 is most highly expressed in adipose tissue, it is also expressed in other cell types such as macrophages and endothelial cells (25–27). FABP4 is widely expressed in the endothelial cells of the microvasculature, with immunoreactivity detected in capillaries and small veins of the heart, lung, kidney, liver, muscle and pancreas (26, 27). Single cell RNA sequencing of 20 mouse organs and tissues revealed FABP4 expression in endothelial cells of 9 of these tissue types, with 72-98% of endothelial cells in liver, kidney, pancreas, adipose tissue, muscle and heart showing positivity for FABP4 expression (28). However, despite this broad expression profile, little is known about the physiological role of endothelial FABP4.

Endothelial FABP4 may play a role in blood vessel growth, as absence of FABP4 in endothelial cells reduces proliferation (27) and siRNA knockdown of endothelial FABP4 decreases angiogenesis in tumors (29). Endothelial FABP4 may also play a role in fatty acid transport into tissues, as siRNA knockdown of FABP4 in human cardiac microvessel endothelial cells results in reduced pioglitazone-mediated fatty acid uptake (30). *In vivo*, FABP4/5 knockout mice have reduced heart, red muscle, and adipose uptake of the fatty acid analog ^125^I-BMIPP (26), though it cannot be ruled out that absence of circulating rather than intracellular endothelial FABP4/5 contributes to this phenotype. Given the widespread expression of FABP4 in endothelial cells, it is important to understand the FABP4 secretory capacity of the endothelium and the physiological and pathophysiological roles of endothelial FABP4 to guide potential translational efforts and strategies.

To investigate the contribution of adipocytes, endothelial cells, and macrophages to the circulating FABP4 pool and the physiological roles of these sources of FABP4, we generated mice with adipocyte, endothelial, myeloid, or whole-body deletion of FABP4. In these studies, we observed that myeloid cells did not contribute to either baseline or stimulated FABP4. Surprisingly, we found that ∼75% of baseline circulating FABP4 in lean mice is derived from endothelial cells, whereas only ∼25% was contributed by adipocytes. In contrast, during lipolysis, adipocytes were the main source of increased circulating FABP4. Unexpectedly, despite near-normal FABP4 responses to lipolysis, mice lacking endothelial FABP4 exhibited reduced lipolysis-induced insulin responses, to the same degree as observed in whole-body FABP4-KO mice. This was in contrast to enhanced glucose-stimulated insulin secretion from isolated islets of endothelial FABP4 deletion mice. These data indicate that endothelial cells are the major contributor to baseline circulating FABP4 and that endothelial-source FABP4 is critical for pancreatic beta cell insulin responses to lipolysis.

## Results

### Development and validation of a tissue-specific FABP4 deletion mouse model

To study the physiological role of different tissue sources of FABP4, we developed FABP4 floxed mice expressing human FABP4. The details of the model development are described in the Methods. We crossed FABP4 floxed mice to Adiponectin-Cre, Tek-Cre, and CMV-Cre mice to generate mice with deletion of FABP4 in adipocytes (Adipo-KO), endothelial cells (Endo-KO), and the whole body (Total-KO), respectively (Figure S1). Since Tek-Cre mouse models are also known to delete from some hematopoietic populations (31) and we have confirmed these observations in peritoneal macrophages (Figure S2), we also crossed FABP4 floxed mice with LyzM-Cre mice to delete FABP4 specifically from the myeloid population (Myeloid-KO), in order to further differentiate the potential contributions of endothelial and macrophage FABP4.

We next validated the deletion of FABP4 in the specific tissues. In western blots of perirenal, mesenteric, and brown (BAT) adipose tissue, we observed near complete deletion of FABP4 protein in Adipo-KO mice (Figure 1A, S3A, S3B). In contrast, Endo-KO adipose tissue showed near-normal FABP4 protein expression. Similarly, immunostaining of perigonadal adipose tissue (PGWAT) and BAT showed absence of FABP4 in adipocytes of Adipo-KO, but not Endo-KO mice (Figure 1B, S3C). It should be noted that Total-KO PGWAT and BAT showed some signal in the immunostaining. Since adipose from FABP4/5-KO mice, which we used as a negative control for the antibody, showed no signal at all (Figure S3D), we surmise that this signal represents some cross-reactivity with FABP5. To determine specific deletion of FABP4 in adipocytes of Adipo-KO mice, we fractionated the PGWAT depot and performed western blots using protein preparations from isolated adipocytes. As in whole adipose tissue, isolated adipocytes of Adipo-KO mice had near complete deletion of FABP4 protein, whereas adipocytes from Endo-KO mice did not differ in FABP4 protein expression from WT adipocytes (Figure 1C).

**Figure 1:**
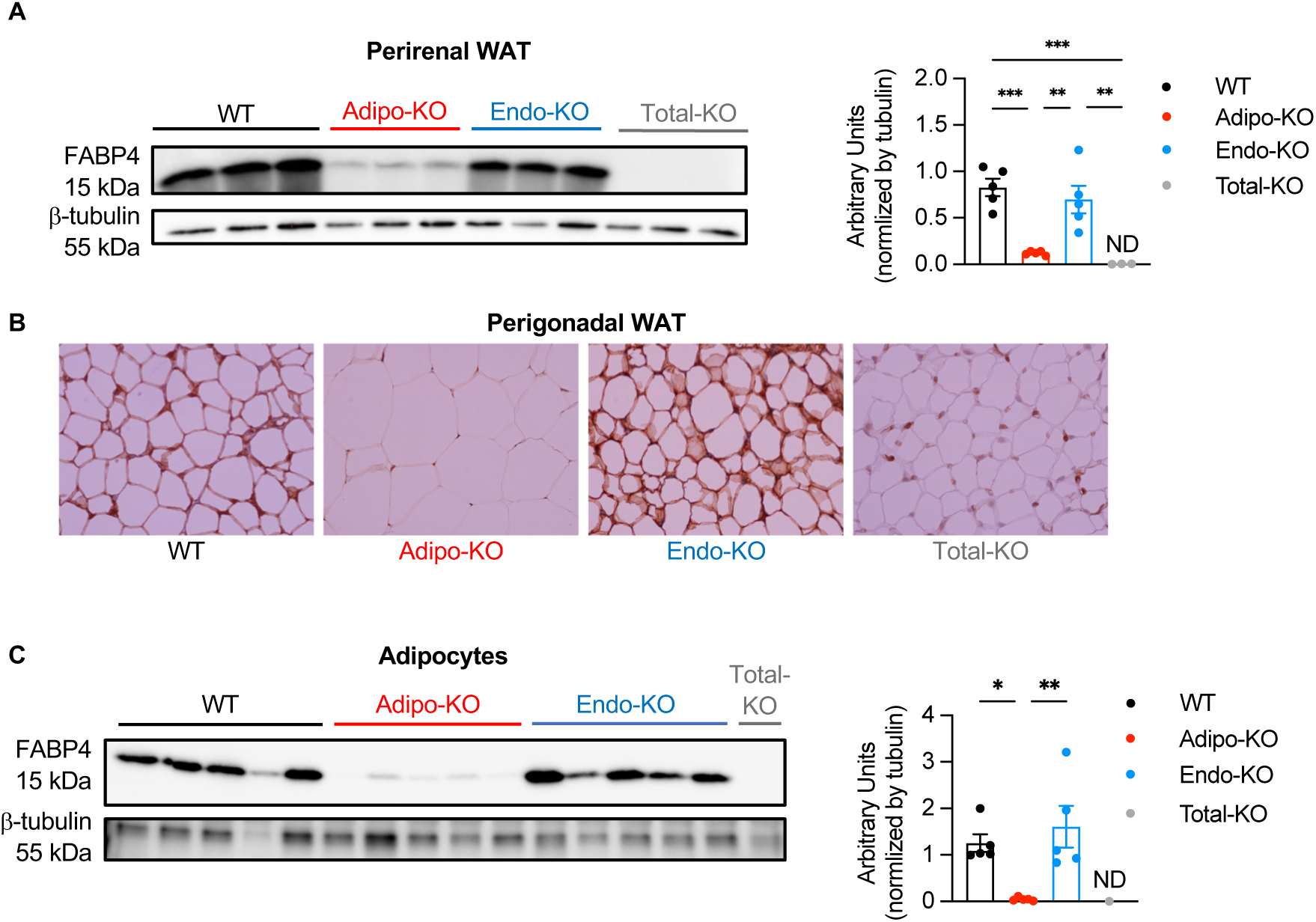
Validation of tissue-specific deletion of FABP4 from adipocytes. (**A**) Left panel: Representative immunoblot of FABP4 expression in perirenal adipose tissue of WT, Adipo-KO, Endo-KO, and Total-KO mice. Right panel: Quantification of FABP4 relative to β-tubulin signal.***p<0.001, **p<0.01. (**B**) FABP4 immunostaining in perigonadal adipose tissue from WT, Adipo-KO, Endo-KO, and Total-KO mice. 40X magnification. (**C**) Left panel: Immunoblot of FABP4 expression in isolated PGWAT adipocytes of WT, Adipo-KO, Endo-KO, and Total-KO mice. Right panel: Quantification of FABP4 relative to β-tubulin signal.*p<0.05, **p<0.01. n=1 for Total-KO. Data were analyzed by one-way ANOVA followed by Tukey’s multiple comparison test and are presented as mean ± SEM. ND = No signal detected.

To confirm deletion of FABP4 from endothelial cells of Endo-KO mice, we isolated endothelial cells from collagenase-digested liver using magnetic beads coated with an antibody against CD31, and performed immunoblots for FABP4 protein. Since CD31 is also expressed on Kupffer cells, we checked the purity of WT endothelial cells isolated with CD31 antibody by co-staining with antibodies against CD31 and the macrophage marker, F4/80, and performed FACS analysis to determine the percentage of endothelial cells versus macrophages (Figure S4A). Of the total pool of cells, 88.5% were positive for CD31 alone and 9.4% were co-stained with anti-CD31 and -F4/80, indicating that 88.5% of the cells were endothelial cells and 9.4% were macrophages. Isolated CD31-positive cells from Endo-KO livers showed no FABP4 signal in western blots, whereas endothelial cells from Adipo-KO livers showed the same FABP4 expression pattern as WT controls (Figure 2A). We also examined FABP4 by immunostaining studies in whole liver tissue. In WT mice, FABP4 positivity was observed in cells lining blood vessels and sinusoids, suggesting FABP4 expression in vascular and sinusoidal endothelial cells or Kupffer cells (Figure 2B). Adipo-KO mice showed similar FABP4 staining patterns to WT mice, whereas Endo-KO mice showed no FABP4 signal in the liver. Western blots of whole liver lysates also showed absence of FABP4 in Endo-KO livers, similar to Total-KO mice (Figure S4B). Since Tek-Cre can delete in some hematopoietic populations (31), and Kupffer cells are also expected to express FABP4, we also performed FABP4 immunostaining in livers of Myeloid-KO mice. Myeloid-KO mice showed FABP4 staining similar to WT mice (Figure 2B), supporting that the absence of FABP4 signal in Endo-KO mice was mainly due to deletion of FABP4 in endothelial cells.

**Figure 2:**
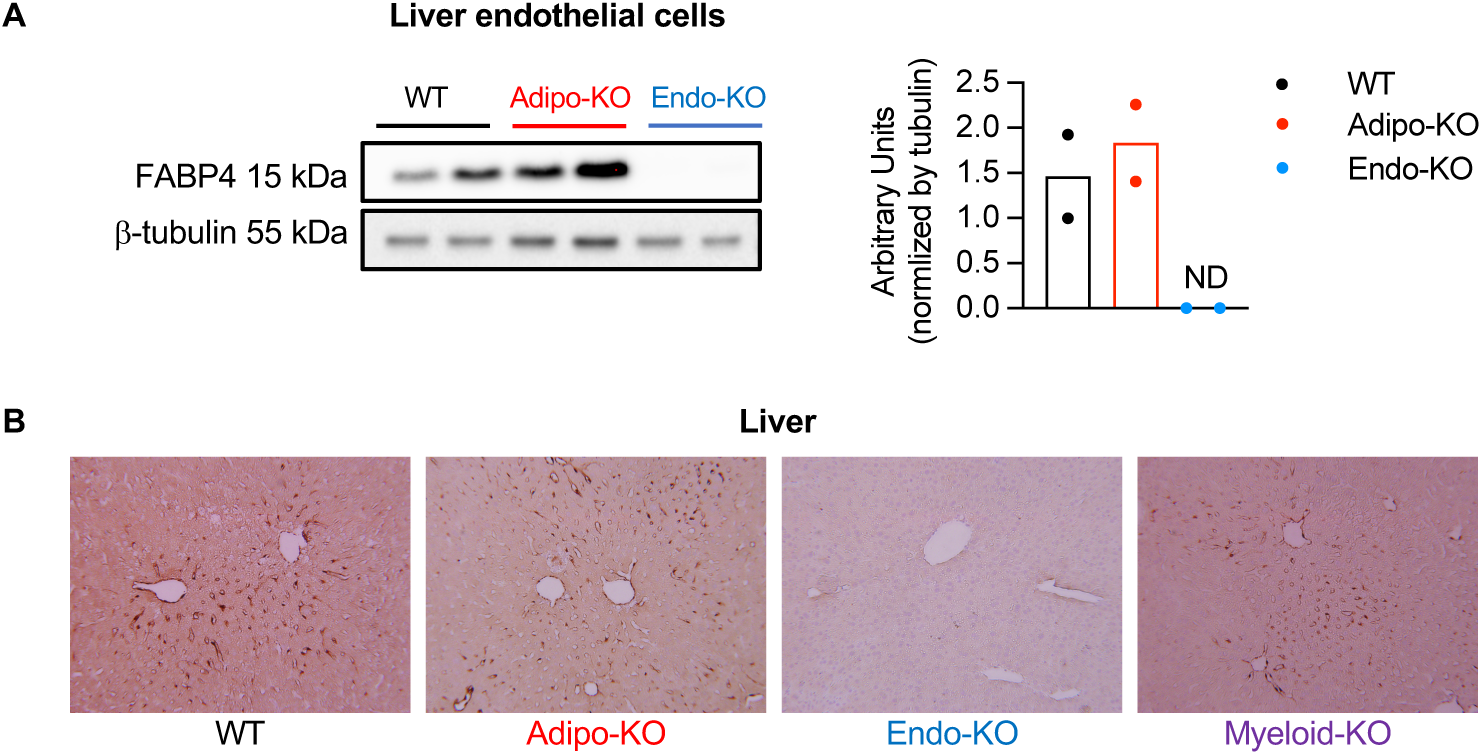
Validation of tissue-specific deletion of FABP4 from endothelial cells. (**A**) Left panel: Immunoblot of FABP4 expression in isolated liver endothelial cells of WT, Adipo-KO, and Endo-KO mice. Right panel: Quantification of FABP4 relative to β-tubulin signal. (**B**) FABP4 immunostaining in liver from WT, Adipo-KO, Endo-KO, and Myeloid-KO mice. 20X magnification. ND = No signal detected.

### Endothelial cells contribute ∼75% of basal circulating FABP4 in lean mice

We and others have shown that adipocytes secrete FABP4 and that circulating FABP4 levels correlate positively with adiposity in mice and humans (12–16). Though FABP4 is expressed in endothelial cells of many tissues and organs (26–28), it is not known whether endothelial cells contribute to circulating FABP4. To examine this important question, we measured circulating FABP4 levels in lean WT, Adipo-KO, and Endo-KO mice. The data shown are a pool of FABP4 levels from multiple experiments in which blood was collected from lean male mice following 6h daytime food withdrawal. To our surprise, Adipo-KO mice only showed ∼25% reduction in baseline circulating FABP4 levels compared to WT controls (Figure 3A). In contrast, Endo-KO mice showed ∼75% decreased basal plasma FABP4 concentrations. We also observed similar results in female mice (Figure S6B). Conversely, Myeloid-KO mice did not show any changes in circulating FABP4 levels (Figure 3B), indicating that these cells do not contribute to FABP4 hormone at measurable levels, and that the reduction in Endo-KO mice was indeed due to absence of FABP4 in endothelial cells. These data are in line with our previous observations in WT mice transplanted with FABP4 knock-out bone marrow, which also did not exhibit alterations in circulating FABP4 (13). Taken together, these data therefore indicate that in lean mice the majority of baseline circulating FABP4 is derived from endothelial cells, rather than adipocytes.

**Figure 3.**
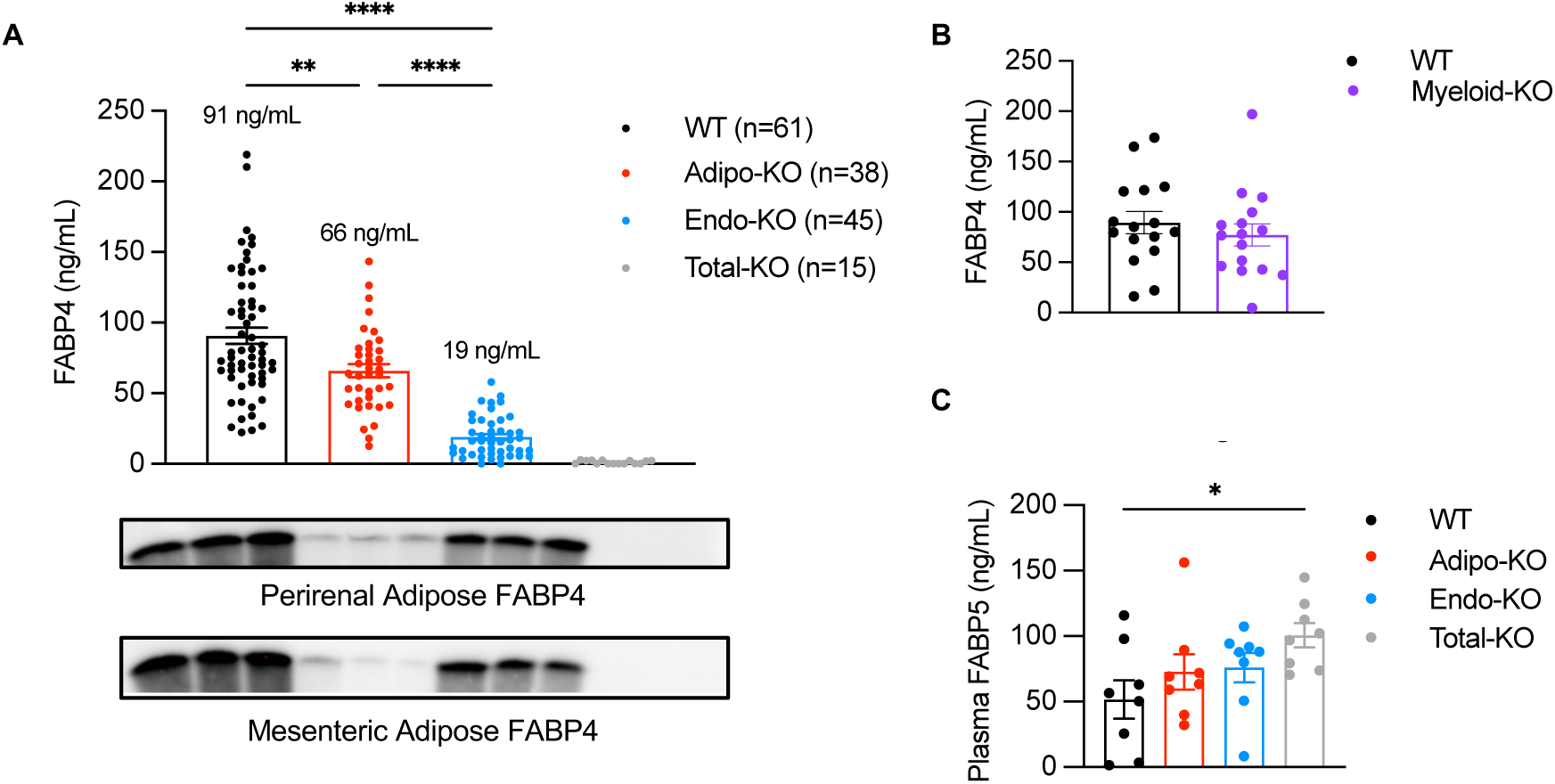
Endothelial cells contribute to ∼75% of basal circulating FABP4. (**A)** Top panel: Plasma FABP4 levels in WT, Adipo-KO, Endo-KO, and Total-KO lean male mice, after 6h daytime food withdrawal. Data are pooled from samples from 9 experiments. ****p<0.0001, **p<0.01. Bottom panel: Immunoblots of perirenal and mesenteric adipose FABP4 protein expression (same images as in Fig 2A and S2A), for comparison with plasma levels. (**B**) Plasma FABP4 levels in 6h WT vs. Myeloid-KO mice. Data are pooled from samples from 2 experiments. (**C**) Plasma FABP5 levels in WT, Adipo-KO, Endo-KO and Total-KO mice. *p<0.05. Data in 3A, 3C, and 3D were analyzed by one-way ANOVA, followed by Tukey’s multiple comparison test. Data in 3B were analyzed by unpaired t-test. Data are presented as mean ± SEM. ND: No signal detected.

In addition to FABP4, adipocytes and endothelial cells also express the closely related isoform, FABP5 (32, 33). Plasma FABP5 levels are nearly doubled in FABP4-KO mice compared to WT controls (13), and adipose or adipocyte levels are at least 7-fold higher (23, 34). In Adipo-KO and Endo-KO mice we did not see significant increases in plasma FABP5 levels compared to WT mice (Figure 3C). Only whole-body FABP4-KO mice showed increased plasma FABP5 compared to WT controls. In whole perirenal adipose tissue there were no differences in FABP5 protein expression between WT, Adipo-KO, and Endo-KO, but unexpectedly Total-KO mice also did not show increased FABP5 (Figure S5), which was seen in earlier conventional total body deletion models (23, 34). Baseline 6h fasting glucose and insulin levels did not differ amongst WT, Adipo-KO, Endo-KO, and Total-KO mice matched for body weight (Table 1). This is not unexpected given that we have previously not observed any differences in these parameters between lean WT and FABP4-KO mice (3).

**Table 1:** Glucose and insulin levels in weight-matched 13-18 week old male WT, Adipo-KO, Endo-KO and Total-KO mice. Blood glucose and insulin levels are after 6h daytime fasting. P=NS. Data are presented as mean ± SEM. N=8/group.

|  | WT | Adipo-KO | Endo-KO | Total-KO |
| --- | --- | --- | --- | --- |
| Body weight (g) | 32.2 ± 0.7 | 32.7 ± 0.9 | 32.4 ± 0.7 | 32.9 ± 0.8 |
| Blood glucose (mg/dL) | 152 ± 7 | 158 ± 5 | 148 ± 4 | 147 ± 6 |
| Insulin (ng/mL) | 0.83 ± 0.11 | 0.83 ± 0.07 | 0.68 ± 0.07 | 0.61 ± 0.10 |

### Lipolysis-driven FABP4 secretion is derived mainly from adipocytes

We have previously shown that lipolysis is a key driver of adipose secretion of FABP4 (13, 14). Stimulation of lipolysis in adipose tissue explants or adipocytes induces lipolysis-dependent increases in FABP4 secretion (14). Similarly, in mice, induction of lipolysis with the β-adrenergic agonist, isoproterenol, or the β3-specific agonist, CL-316,243, rapidly induces increases in circulating FABP4 (13). To determine the cellular source of the lipolysis-induced increases in circulating FABP4, we compared FABP4 secretion in WT, Adipo-KO and Endo-KO mice following isoproterenol-induced lipolysis. Despite only mildly reduced basal plasma FABP4 levels, male Adipo-KO mice showed significantly diminished FABP4 secretory responses to isoproterenol compared to WT controls (Figures 4A,S6A). This corresponded to a 62% decrease in the area under the curve (AUC) of the Adipo-KO response compared to WT controls. In contrast, despite a 75% reduction in baseline FABP4 levels, Endo-KO mice had FABP4 responses much closer to WT mice. A similar pattern of FABP4 secretion in response to β-adrenergic stimulation was observed in female mice (Figure S6C). Non-esterified fatty acid (NEFA) responses to induction of lipolysis were decreased at the 30 min time point in Total-KO mice compared to WT (Figure 4B). Glycerol responses to isoproterenol-induced lipolysis were also mildly decreased in Total-KO and Adipo-KO mice vs. WT controls (Figure 4C). These data are in line with previous observations of mildly decreased glycerol responses in whole-body FABP4-KO mice (23, 34), and show that absence of adipocyte FABP4 may underlie this phenotype.

**Figure 4.**
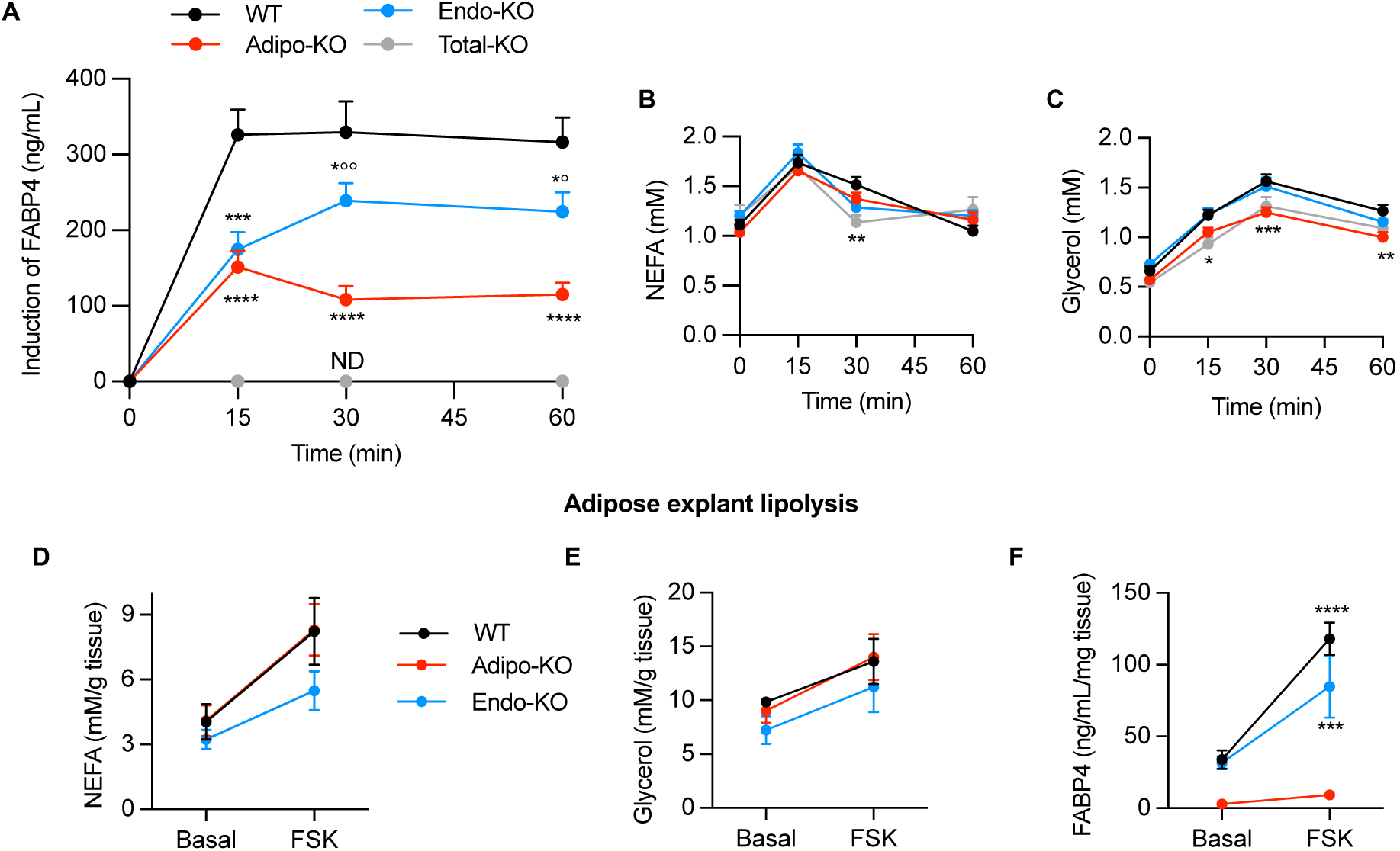
Lipolysis-driven FABP4 secretion is primarily from adipocytes. (**A**) Plasma FABP4, (**B**) Non-esterified fatty acid (NEFA), and (**C**) glycerol responses to 10mg/kg isoproterenol-induced lipolysis in WT, Adipo-KO, Endo-KO, and Total-KO male mice. FABP4 responses are presented as induction over baseline. Data are pooled from 6 separate experiments. WT n=52, Adipo-KO n=39, Endo-KO n=35, Total-KO n=8 for Fig 4A, n=15 for Figs 4B, 4C. ****p<0.0001, ***p<0.001, **p<0.01, *p<0.05 vs. WT; °°p<0.01 vs. Adipo-KO, °p<0.05 vs. Adipo-KO. (**D**) NEFA, (**E**) glycerol, and (**F**) FABP4 responses to forskolin (FSK)-induced lipolysis in perigonadal adipose explants from male WT, Adipo-KO, and Endo-KO mice. Data are normalized to amount of adipose tissue per culture well. n=4 mice/group, 3 replicates per mouse. ****p<0.0001, ***p<0.001 vs. Adipo-KO. Data were analyzed by repeated measures two-way ANOVA followed by Tukey’s multiple comparison test and are presented as mean ± SEM. ND: No signal detected.

Notably, adipocyte deletion of FABP4 did not completely abrogate the FABP4 response to isoproterenol, and likewise, Endo-KO mice still had a 33% reduction in the AUC of the FABP4 response. To rule out whether the small induction of FABP4 in Adipo-KO mice may have been due to residual adipose secretion of FABP4, which could occur if there was incomplete deletion in adipocytes, we performed forskolin (FSK)-induced lipolysis in adipose explants from WT, Adipo-KO, and Endo-KO mice. There was no effect of adipose or endothelial deletion of FABP4 on explant NEFA and glycerol responses to FSK (Figures 4D,E). Induction of lipolysis in explants led to robust and similar increases in FABP4 from WT and Endo-KO explants (Figure 4F). Importantly, Adipo-KO explants showed no baseline or FSK-induced FABP4 response. These data suggest that a fraction of the FABP4 secretion observed in Adipo-KO mice *in vivo* may in fact be derived from the endothelial cell population.

Since Tek-Cre also deletes from myeloid cells we performed lipolysis in WT vs. Myeloid-KO mice to determine whether the mild reduction in FABP4 responses in Endo-KO mice was due to absence of macrophage FABP4. FABP4 responses in Myeloid-KO mice were identical to WT controls (Figure 5A), indicating that the mildly reduced FABP4 response in Endo-KO mice was indeed due to absence of endothelial FABP4. We next confirmed whether the small induction of FABP4 observed in Adipo-KO mice was due to secretion from endothelial cells. To do this we crossed Adipo-KO mice with Endo-KO mice to delete FABP4 in both cell types. Mice with deletion in both depots had nearly undetectable FABP4 responses to lipolysis (Figure 5B), indicating that the residual induction of FABP4 secretion in Adipo-KO mice was indeed from endothelial cells. Thus, while adipocytes supply the majority of the FABP4 response to lipolytic stimulation, we now show that the endothelium may also contribute a minor fraction *in vivo*. As endothelial cells from Adipo-KO mice do not express increased amounts of FABP4 compared to WT endothelial cells (Figure 2A), the endothelial FABP4 response to lipolysis in Adipo-KO mice does not appear to be a compensatory response to lack of adipocyte FABP4.

**Figure 5.**
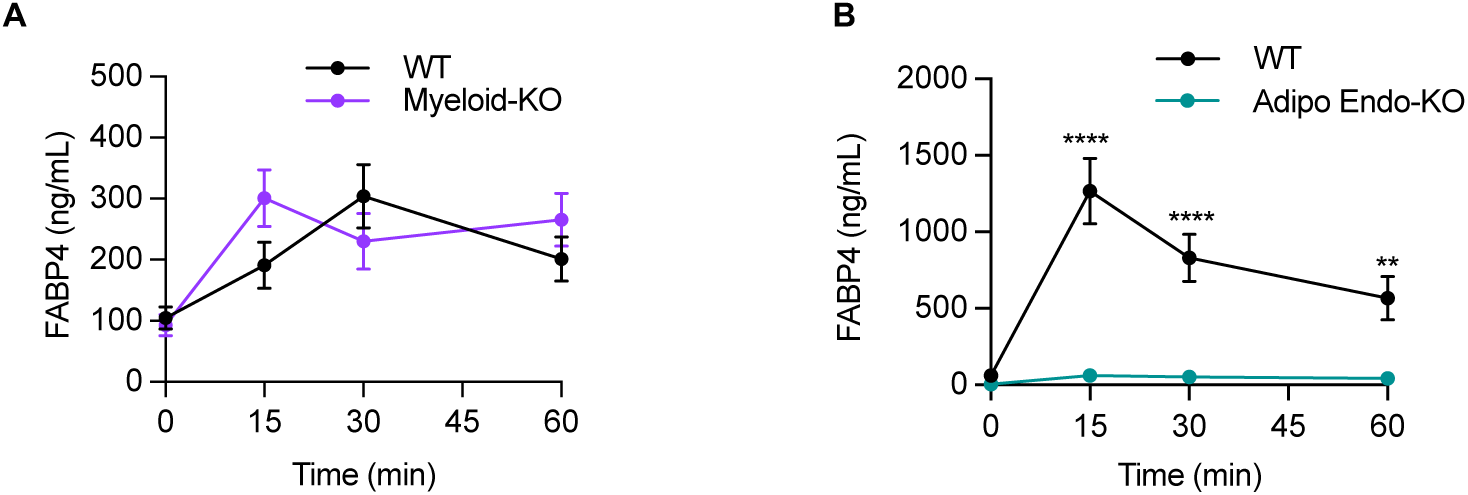
Lipolysis-driven insulin secretion in Myeloid-KO and mice with deletion of FABP4 in adipocytes and endothelial cells. (**A**) Plasma FABP4 responses to 10mg/kg isoproterenol-induced lipolysis in WT and Myeloid FABP4-KO mice, n=8/group, and (**B**) in WT vs. mice with deletion of FABP4 in both adipocytes and endothelial cells (Adipo Endo-KO). n=6/group. ****p<0.0001, **p<0.01. Data were analyzed by repeated measures two-way ANOVA followed by Tukey’s multiple comparison test and are presented as mean ± SEM.

### Regulation of endothelial FABP4 secretion differs from adipocytes

To examine the FABP4 secretory capacity of endothelial cells we cultured isolated endothelial cells (using the CD31 affinity method described above) pooled from liver, heart, and lungs of WT, Adipo-KO, and Endo-KO mice and collected 4-hour conditioned media for assessment of FABP4 levels. WT and Adipo-KO endothelial cells both secreted FABP4 in similar amounts, providing direct evidence that endothelial cells secrete FABP4 (Figure 6A). As expected, Endo-KO cells showed no detectable secreted FABP4 protein in their media. Deletion of FABP4 in endothelial cells in Endo-KO mice did not lead to a compensatory upregulation of FABP5, as FABP5 levels in Endo-KO liver endothelial cell conditioned media and lysates did not differ from WT endothelial cells (Figures 6B,C).

**Figure 6.**
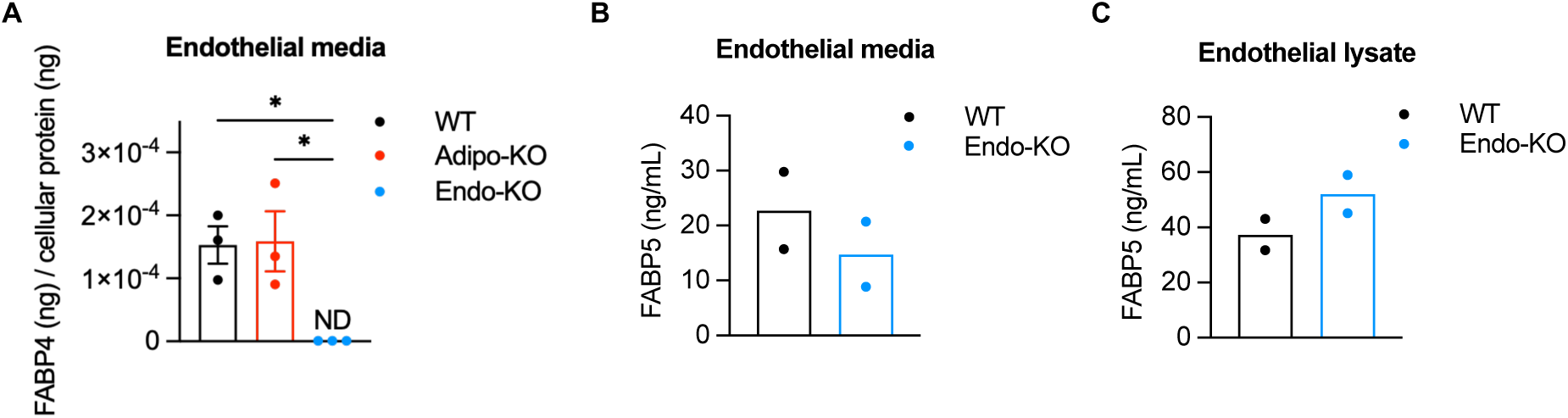
Endothelial cells secrete FABP4. (**A**) 4-hour conditioned media FABP4 levels from CD31-isolated liver, heart, and lung endothelial cells of WT, Adipo--KO, and Endo-KO mice. FABP4 levels are normalized to total cellular protein levels. *p<0.05. (**B**) FABP5 levels in CD31-isolated liver endothelial cell 24-hour conditioned media and **(C**) lysates of WT and Endo-KO mice. Data in 6A were analyzed by one-way ANOVA, followed by Tukey’s multiple comparison test and are presented as mean ± SEM. ND: No signal detected.

To further investigate the potential regulation of FABP4 secretion by endothelial cells, we performed studies in human umbilical vein endothelial cells (HUVECs). We first tested FABP4 expression at different stages of HUVEC culture, as it is known that HUVECs grown to reach the tightly packed monolayer “cobblestone” stage more closely resemble the morphology and phenotype of endothelial cells *in vivo* (35). Protein levels of FABP4 in HUVEC lysates were maximal at day 14 post-seeding, after the cells had reached the cobblestone phenotype (Figures 7A, S7A). As was shown in isolated primary endothelial cells, HUVECs exhibited secretion of FABP4 into the media, with increased secretion beginning on day 7 (Figure 7B). Day 7 HUVECs showed a steady increase in media FABP4 released over 20 hours, while media lactate dehydrogenase (LDH) levels remained unchanged (Figure 7C), indicating that HUVECs were, in fact, secreting FABP4 rather than releasing FABP4 due to cell death.

**Figure 7.**
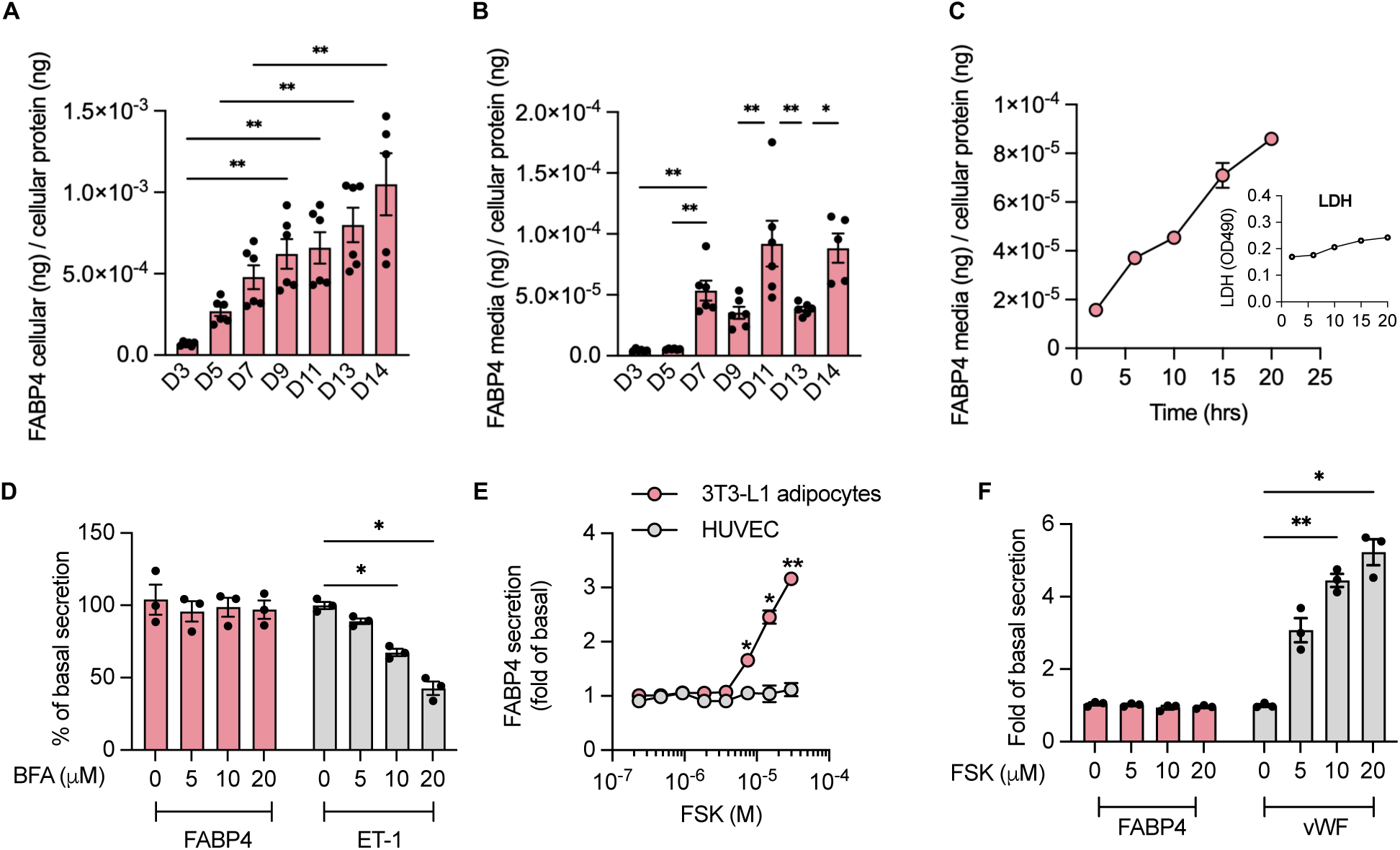
Endothelial FABP4 secretion and is differentially regulated from adipocyte FABP4 secretion. (**A**) FABP4 in HUVEC lysates normalized to total cellular protein (**B**) and in 5-hour conditioned media normalized to total cellular protein at days 3 through 14 post-seeding. Pool of 2 experiments. *p<0.05, **p<0.01. (**C**) Time-course of cumulative FABP4 levels in media of day 7 HUVECs. n=4/time point. Inset: Media lactate dehydrogenase (LDH) levels during the same time course. n=4. (**D**) Effects of increasing doses of the ER-Golgi pathway inhibitor, brefeldin A (BFA) on FABP4 and endothelin-1 (ET-1) secretion from day 11 HUVECs. n=3 per BFA dose. *p<0.05. (**E**) Effects of increasing doses of forskolin (FSK) on FABP4 secretion in HUVECs vs. 3T3-L1 adipocytes. n=3 per FSK dose. *p<0.05, **p<0.01 vs. HUVEC. (**F**) Effects of increasing doses of FSK on FABP4 and Von Willibrand Factor (vWF) secretion from day 11 HUVECs. n=3 per FSK dose. *p<0.05, **p<0.01. 7A-C were analyzed by one-way ANOVA followed by Tukey’s multiple comparison test. 7D-F were analyzed by repeated measures 2-way ANOVA with Geisser-Greenhouse correction, followed by Dunnett’s (D,F) or Sidak’s (E) multiple comparison test. Data are presented as mean ± SEM.

We then tested how endothelial FABP4 secretion compares with adipocyte FABP4 secretion in cobblestone HUVECs at day 11. Adipocyte FABP4 is not secreted via the classical endoplasmic reticulum (ER)-Golgi pathway (14, 15, 36), but is mostly released from secretory lysosomes (15). To determine which pathway mediates endothelial FABP4 secretion, we treated HUVECs with the ER-Golgi pathway inhibitors, brefeldin-A or monensin. HUVEC FABP4 secretion was not inhibited by either agent (Figure 7D, Supplemental Table 1), indicating that, like adipocytes, endothelial FABP4 secretion does not occur through the classical ER-Golgi pathway. Secretion of endothelin-1, a protein secreted via the ER-Golgi pathway (37), was inhibited by brefeldin A (Figure 7D), indicating that the ER-Golgi secretory mechanism was functional in our HUVEC culture. We also tested the effect of the lysosomal secretion inhibitors chloroquine and ammonium chloride. Whereas chloroquine and ammonium chloride inhibit isoproterenol or FSK-induced FABP4 release by adipocytes (15), they did not inhibit FABP4 secretion by HUVECs (Supplemental Table 1). Increased intracellular calcium by treatment with ionomycin has also been shown to induce FABP4 release by human adipocytes (36). Here we treated HUVECs with histamine, which increases intracellular Ca^2+^, but did not see induction of FABP4 secretion (Supplemental Table 1). Taken together, our data show that secretion of FABP4 by endothelial cells is not stimulated by the same mechanisms as in adipocytes.

We next tested the effects of the lipolytic agents, FSK, IBMX, isoproterenol, and CL-316,243 on HUVEC FABP4 secretion. FSK induced robust secretion of FABP4 by 3T3-L1 adipocytes but had no effect on HUVEC FABP4 secretion (Figure 7E). Similarly, IBMX, isoproterenol, and the β3-adrenergic receptor-specific agonist, CL-316,243, had no effect to induce HUVEC FABP4 secretion (Supplemental Table 1). This is in contrast to the effect of isoproterenol to induce endothelial FABP4 secretion *in vivo*. The lack of responsiveness to FSK was not due to suboptimal growing conditions of the HUVEC cultures, as HUVEC secretion of von Willebrand Factor (vWF) responded robustly to FSK treatment (Figure 7F). This difference between *in vivo* and *in vitro* responses to cAMP-mediated stimulation indicates that there may be additional factors required for endothelial FABP4 secretion *in vivo* or that FABP4 may be coming from a specialized endothelial subpopulation that is not represented in these culture systems.

### Lipolysis-driven insulin secretion is blunted in mice lacking endothelial FABP4

In response to β-adrenergic induction of lipolysis, FABP4-KO mice have markedly blunted insulin responses, suggesting that FABP4 deficiency alters islet beta cell function during acute lipolysis (23). To examine whether this phenotype is due to absence of FABP4 in adipocytes or endothelial cells, we measured plasma insulin responses to isoproterenol-induced lipolysis. Surprisingly, despite the markedly blunted FABP4 induction in response to lipolysis, Adipo-KO mice did not show any significant reduction in insulin secretion (Figure 8A). There were also no changes in insulin responses in Myeloid-KO mice (Figure 8B). In contrast, the AUC of insulin responses in Endo-KO mice was reduced by 62% reduced compared to WT mice, to a similar degree displayed by Total-KO mice. These data suggest that endothelial-source FABP4 is a key requirement for lipolysis-driven acute induction of insulin secretion.

**Figure 8.**
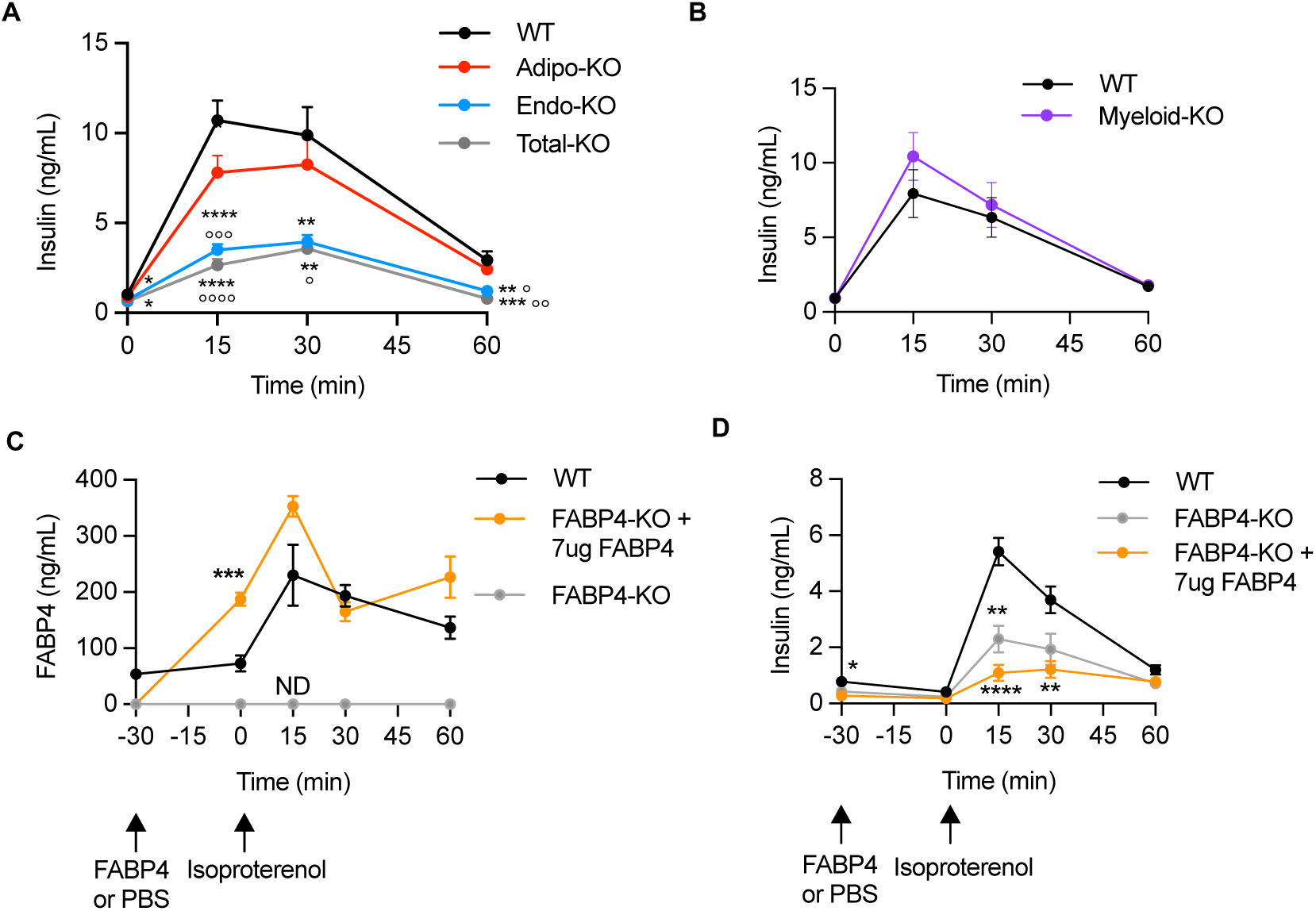
Lipolysis-driven insulin secretion is blunted in mice lacking endothelial, but not adipocyte FABP4. (**A**) Plasma insulin responses to 10mg/kg isoproterenol-induced lipolysis in WT, Adipo-KO, Endo-KO, and Total-KO mice. Data are pooled from 6 experiments. ****p<0.0001, ***p<0.001, **p<0.01, *p<0.05 vs. WT. °°°°p<0.0001, °°°p<0.001, °°p<0.01, °p<0.05 vs. Adipo-KO. WT, n=51; Adipo-KO, n=39; Endo-KO, n=34; Total-KO, n=15. (**B**) Plasma insulin responses to 10mg/kg isoproterenol-induced lipolysis in WT vs. Myeloid-KO mice. n=8/group. (**C**) Plasma FAPB4 and (**D**) insulin responses in WT and FABP4-KO mice injected with PBS or 7ug of FABP4 prior to induction of lipolysis with 10mg/kg isoproterenol. n=8/group. ***p<0.0005, **p<0.001, *p<0.05 vs. WT. Data were analyzed by repeated measures 2-way ANOVA with Geisser-Greenhouse correction, followed by Sidak’s (C) or Tukey’s (A,D) multiple comparison test. Data are presented as mean ± SEM. ND: No signal detected.

We sought to determine the underlying causes of the decreased insulin responses in Endo-KO mice. One possible contribution is the low basal FABP4 levels in Endo-KO mice. To investigate this question, we administered FABP4-KO mice with recombinant FABP4 by intraperitoneal injection 30 minutes prior to induction of lipolysis. Despite reaching comparable FABP4 levels to wildtype controls (Figure 8C), insulin responses were not restored by FABP4 administration, and even tended to be reduced further (Figure 8D), an effect we have previously observed with *in vivo* FABP4 administration (24). These data suggest a few possibilities - that deficiency of FABP4 within endothelial cells may underlie the lack of lipolysis-associated insulin secretion, that endothelial-source FABP4 functions differently from adipocyte-source FABP4, a factor which is not captured by the use of recombinant protein, or that chronic low circulating FABP4 levels in Endo-KO and Total-KO mice may alter beta cell insulin responsiveness during lipolysis.

### Deficiency of endothelial FABP4 alters the insulin secretory response

The pancreas is highly vascularized, and endothelial cells are known to secrete a variety of factors that influence beta cell function (38). To examine the potential role for endothelial FABP4 in insulin secretion from islets we performed immunostaining for FABP4 and insulin on pancreatic sections and examined the *ex vivo* insulin secretion phenotype of WT, Adipo-KO and Endo-KO mouse islets. In our earlier studies, we demonstrated that no FABP4 expression is present in isolated islets or beta cell lines (23, 24). Consistently, pancreata from WT and Adipo-KO mice showed minimal FABP4 immunoreactivity in islets (Figure 9A). However, there was FABP4 staining in the vasculature of the exocrine pancreas. This FABP4 signal was completely lost in the entire pancreas of Endo-KO mice but was observed in Myeloid-KO mice, suggesting that the FABP4-positive cells were primarily endothelial cells. Recent evidence supports an integrated capillary network of bidirectional blood flow between the exocrine and endocrine pancreas (39). Together these data indicate that FABP4 generated from endothelial source may be the effector on beta cells.

**Figure 9.**
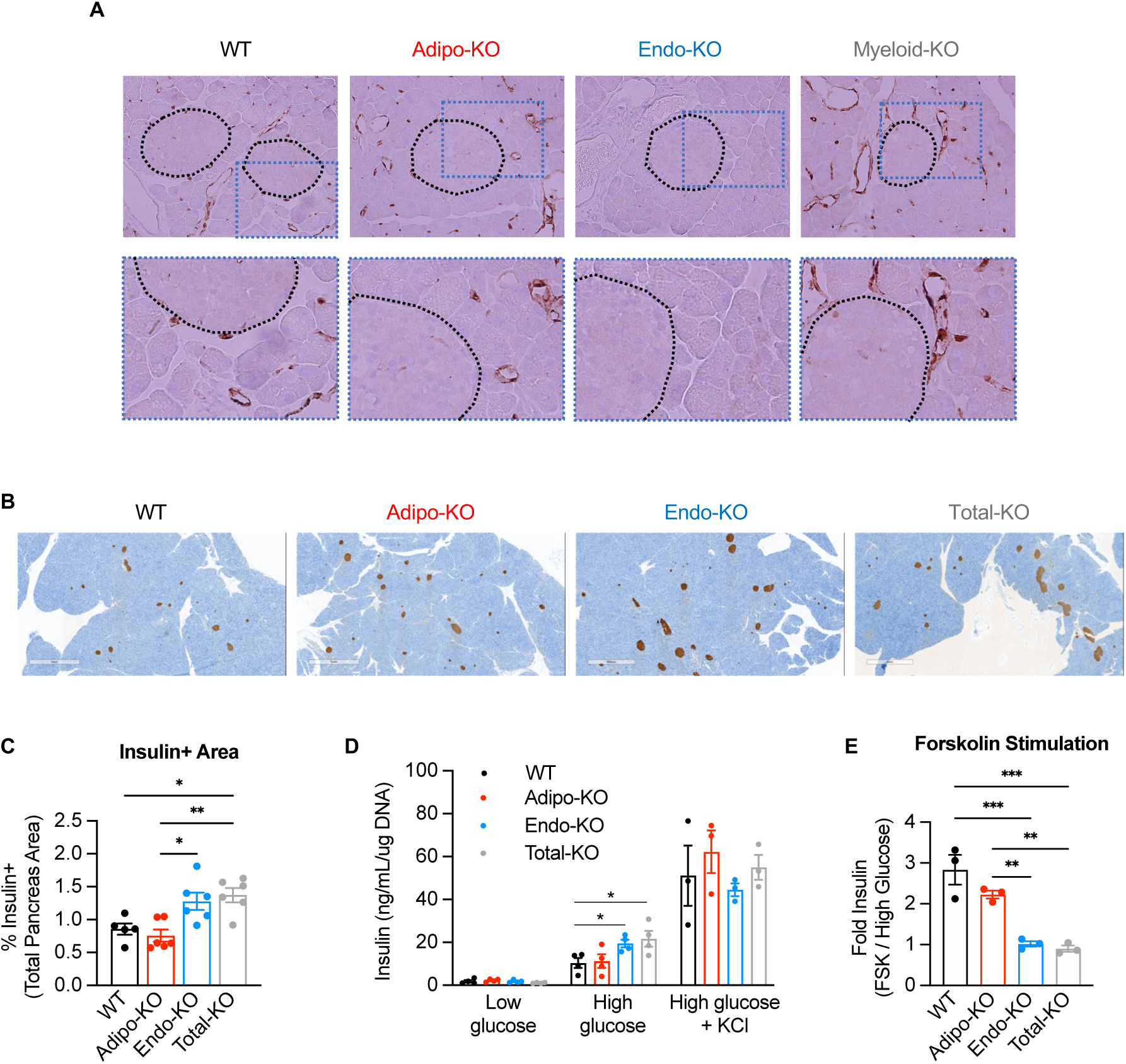
Pancreas endothelial cells express FAPP4 and Endo-KO islets show altered regulation of insulin secretion. (**A**) Upper panel: FABP4 immunostaining of pancreas of WT, Adipo-KO, Endo-KO and Myeloid-KO mice. Islets are encircled by black dotted lines. 40X magnification. Lower panel: 2X enlargement of boxed area in upper panel. (**B**) Insulin immunostaining of pancreas from WT, Adipo-KO, Endo-KO, and Total-KO mice, 40X magnification, and (**C**) quantification of insulin-positive area. (**D**) Insulin secretion from isolated islets of WT, Adipo-KO, Endo-KO and Total-KO mice in response to low glucose (2.8mM), high glucose (16.7mM), high glucose + KCl (30mM). *p<0.05. (**E**) Fold-increase in insulin secretion induced by HG + FSK (10uM) over HG. Data in D were analyzed by unpaired t-test. Data in C and E were analyzed by one-way ANOVA followed by Tukey’s multiple comparison test. Data are presented as mean ± SEM.

Despite the reduced lipolysis-induced insulin secretion in Endo-KO and Total-KO mice, staining of pancreatic sections revealed increases in insulin-positive area in these mice compared to WT and Adipo-KO mice (Figures 9B,C), similar to our previous observation of increased beta cell mass in FABP4-KO mice (24). In contrast to *in vivo* insulin responses during lipolysis, primary islets isolated from Endo-KO and Total-KO mice showed increased glucose-stimulated insulin secretion (GSIS) compared to both Adipo-KO and WT islets (Figure 9D). This was not due to a general increase in insulin secretion, as addition of the secretagogue KCl showed no difference in the maximal secretory capacity between all groups. These findings are supported by our previous results in whole-body FABP4-KO mice, where the response to the insulin secretagogue L-arginine was intact (23) and islets from these animals exhibited a superior GSIS (24). In contrast to high glucose, however, further induction of insulin secretion by increasing intracellular cAMP with FSK was reduced in Endo-KO and Total-KO islets compared to WT and Adipo-KO (Figure 9E), mirroring the reduced insulin response to isoproterenol-induced lipolysis *in vivo*. Taken together, our data demonstrate that islets from Endo-KO and Total-KO mice show reduced cAMP-mediated, but enhanced glucose-induced insulin secretion.

## Discussion

Since the discovery that adipocytes secrete FABP4 and that circulating FABP4 is elevated in obesity (12, 13), it has generally been presumed that adipocytes are the major source of plasma FABP4 and are responsible for its hormonal effects. Here, unexpectedly, we report that in lean mice under basal unstimulated conditions adipocytes only contribute ∼25% of steady-state circulating FABP4, and instead, endothelial cells are the predominant source, contributing to ∼75% of baseline plasma FABP4. In contrast, in response to β-adrenergic-induced lipolysis, adipocytes are the major source of lipolysis-driven increases in circulating FABP4. Surprisingly, despite the significantly diminished FABP4 responses in mice with adipocyte-specific FABP4 deficiency, these animals have essentially intact acute insulin responses to lipolysis, whereas mice lacking endothelial FABP4 have similarly diminished insulin responses to lipolysis as the whole-body FABP4-KO mice, despite only mildly reduced FABP4 secretion. We demonstrate for the first time that the endothelium is the major source of baseline circulating FABP4, and that endothelial FABP4 is a key requirement for lipolysis-induced insulin secretion.

Our data is the first evidence that endothelial cells contribute to circulating FABP4 *in vivo,* predominantly under basal conditions, but also to a minor extent under stimulated conditions. Our examination of FABP4 secretion in cultured endothelial cells indicates that, like in adipocytes, secretion does not occur through the classical ER-Golgi pathway (13, 15). However, unlike adipocytes, at least *in vitro*, endothelial FABP4 secretion appears to be constitutive, rather than inducible. Endothelial FABP4 secretion is not regulated in a similar manner to adipocytes and a major difference is that, at least *in vitro*, endothelial cells do not secrete FABP4 in response to β-adrenergic stimuli and increases in intracellular cAMP. Since it is established in other contexts that endothelial cells can respond to isoproterenol, for example by increasing eNOS activity via β2-adrenergic receptor activation (40), this is unlikely to be the result of a general lack of β-adrenergic signaling machinery in these cells. Rather, these data suggest that regulated secretion of FABP4 may not be due to a direct effect of isoproterenol on endothelial cells, or that *in vitro* culture conditions are not replicating the *in vivo* conditions required for endothelial cell FABP4 secretion in response to β-adrenergic or other stimuli. Regardless, endothelial cells and adipocytes differ in their responses to β-adrenergic stimulation in terms of FABP4 secretion *in vitro* and further studies are warranted to explore the potential signals that may induce secretion in whole animals.

The differential cellular sourcing of baseline versus induced FABP4 secretion may have significant implications for understanding the pathological versus physiological effects of circulating FABP4. It is well-established that FABP4-deficient mice are protected against obesity-induced hyperinsulinemia, insulin resistance and glucose intolerance (3, 5). It is also well-established that basal lipolysis is constitutively increased in obesity (41), and therefore one possible mechanism for protection of FABP4-deficient mice from insulin resistance and its consequences may be decreased lipolysis-mediated insulin secretion, which is significantly reduced in these animals (23). Recent evidence supports a model where prevention of hyper-secretion of insulin and hyperinsulinemia may be critical for alleviation of insulin resistance and protecting beta cell functionality and survival (42). Free fatty acids (FFAs) possess an important role in this regulatory circuit, as they are critical for both glucose- non-glucose-induced insulin secretion (43–45). FFA-mediated stimulation of insulin secretion is known to occur through activation of the fatty acid receptor FFAR1 (aka GPR40) on beta cells (46–48). Here, whole-body and endothelial FABP4-KO mice show markedly blunted insulin responses to β-adrenergic-induced lipolysis despite only minimal or no reductions in NEFA and glycerol responses and improved glucose-stimulated insulin secretion in isolated islets. These data indicate that in addition to FFAs, FABP4 specifically from endothelial cells is a key regulator for lipolysis-mediated insulin secretion. Moreover, endothelial FABP4 promotion of lipolysis-mediated insulin secretion may be detrimental for beta cell health and function, as hyperinsulinemia is a driver for the development of insulin resistance (49).

Our data demonstrate that neither the total body, nor the endothelial-specific deletion of FABP4, results in defective insulin secretion or decreased islet insulin content. In fact, both islet insulin content and insulin responses to glucose were increased in these models, supporting the suppressive role of FABP4 on pancreatic beta cell mass and the proper function of islets (24). In contrast, islets isolated from either of these models show a blunted insulin response to FSK stimulation, suggesting that absence of endothelial FABP4 decreases cAMP-mediated amplification of insulin secretion. cAMP is not known to be induced by FFA through FFAR1/GPR40, which is a Gχξq receptor whose activation by fatty acids induces insulin secretion via the IP3 and DAG pathway (50, 51). GPR40 knockout mice only show partially decreased insulin responses to β-adrenergic-induced lipolysis (52), also suggesting that other signaling pathways are involved in lipolysis-mediated insulin secretion. Though FABP4 has capability to bind fatty acids, our data suggest that endothelial FABP4 may at least in part be acting independently of FFAR1/GPR40 signaling to modulate insulin secretion via a cAMP-dependent mechanism.

In summary, we show that endothelial cells are the major contributor to baseline circulating FABP4, whereas adipocytes are primarily responsible for lipolysis-driven increases in FABP4. Surprisingly, however, endothelial FABP4 rather than adipocyte FABP4 is the critical requirement for lipolysis-driven insulin secretion. Our observation of increased islet mass and changes in islet insulin secretion that persisted *ex vivo* in mice lacking endothelial FABP4 suggests a potential developmental role for endothelial FABP4 in beta cell proliferation and function. This is intriguing, given that FABP4-KO mice show decreased endothelial responsiveness to the proliferative effects of vascular endothelial growth factor (VEGF) (53), and VEGF has an important role in pancreatic vascular development, which in turn influences beta cell function (54). Alternatively, endothelial FABP4 may have an acute effect on islets to stimulate insulin secretion. The effect of endothelium to secrete FABP4 and modulate insulin secretion indicates a role for the endothelium as an endocrine organ and a potential key regulator of metabolism. In future studies, it will be important to determine the cellular sources of FABP4 and impact on the pathogenesis of conditions such as cancer, obesity, and cardiovascular disease, where its circulating levels are regulated and known to be critical for systemic disease.

## Methods

### Generation of FABP4 flox mice and tissue-specific deletion models

Studies were performed in humanized FABP4 flox mice on C57BL/6 background generated for the Hotamışlıgil lab by GenOway. The *FABP4* gene is located on chromosome 3 and consists of 4 exons encoding a 132 amino acid, 15kDa protein. A targeting strategy was developed based on analysis of the FABP4 gene organization and the predicted functional protein structure. The murine FABP4 sequence was first converted to human FABP4 by introducing point mutations to convert the 11 amino acids in exons 2 and 3 that differ between mouse and human FABP4 to human FABP4 sequences (Figure 10A). This will allow for future studies to develop targeting therapies against human FABP4. A single loxP site was inserted within intron 1 upstream of exon 2 and a FRT-neomycin-FRT-loxP cassette was inserted at the 3’ end of exon 2 (Figure 10B). The targeting construct was electroporated into C57BL/6 embryonic stem (ES) cells. The positive ES cell clones were injected into blastocysts of albino C57BL/6J mice (C57BL/6J-Tyrc^-2J^/J), which were implanted into pseudo-pregnant females. Highly chimeric male pups were bred to whole-body Flp recombinase-expressing mice to excise neomycin selection cassette. The resulting FABP4 flox mice were crossed to Adiponectin-Cre (B6.FVB-Tg(Adipoq-cre)1Evdr/J, Jackson Labs stock #028020)(55), Tek-Cre (B6.Cg-Tg(Tek-cre)1Ywa/J, Jackson Labs, stock # 008863)(56), LyzM-Cre (B6.129P2-*Lyz2^tm1(cre)Ifo^*/J, Jackson Labs, stock #004781)(57), and CMV-Cre (B6.C-Tg(CMV-cre)1Cgn/J, Jackson Labs, stock #006054)(58), to generate mice with deletion of FABP4 in adipocytes (Adipo-KO), endothelial cells (Endo-KO), myeloid cells (Myeloid-KO) and the whole body (Total-KO), respectively (Figure S1). Since FABP4 flox CMV-Cre-positive mice have germline expression of the truncated FABP4, the Cre recombinase was bred out of these mice by crossing to wildtypes and retaining mice with only the truncated FABP4 gene to cross with each other to generate homozygous FABP4 flox-KO (Total-KO). Initially, FABP4 flox mice were genotyped by in-house by PCR from tail biopsies (fwd: 45835-Flp-HCA1 5’ GAAGGTGAGGAACAAGGAGCTGATGC 3’; rev: 45836-Flp-HCA1 5’ AGGTGGGCACGGTAATGTTATGGTG 3’, wildtype 333bp, knock-in 453bp). Subsequently, genotyping for FABP4 flox, FABP4 flox-KO, and Cre recombinase was commercially outsourced and performed from tail biopsies or ear punches using real time PCR with specific probes designed for each gene (Transnetyx, Cordova, TN). FABP4 flox mice expressing both Adiponectin and Tek-Cre were genotyped with Cre probes that were specific for Adiponectin-Cre and Tek-Cre. Mice were studied at 8-18 weeks of age and were maintained on a 12-hour-light/12-hour-dark cycle in the Harvard T.H. Chan School of Public Health pathogen-free barrier facility with free access to water and to a standard laboratory chow diet (PicoLab Mouse Diet 20 #5058, LabDiet).

**Figure 10:**
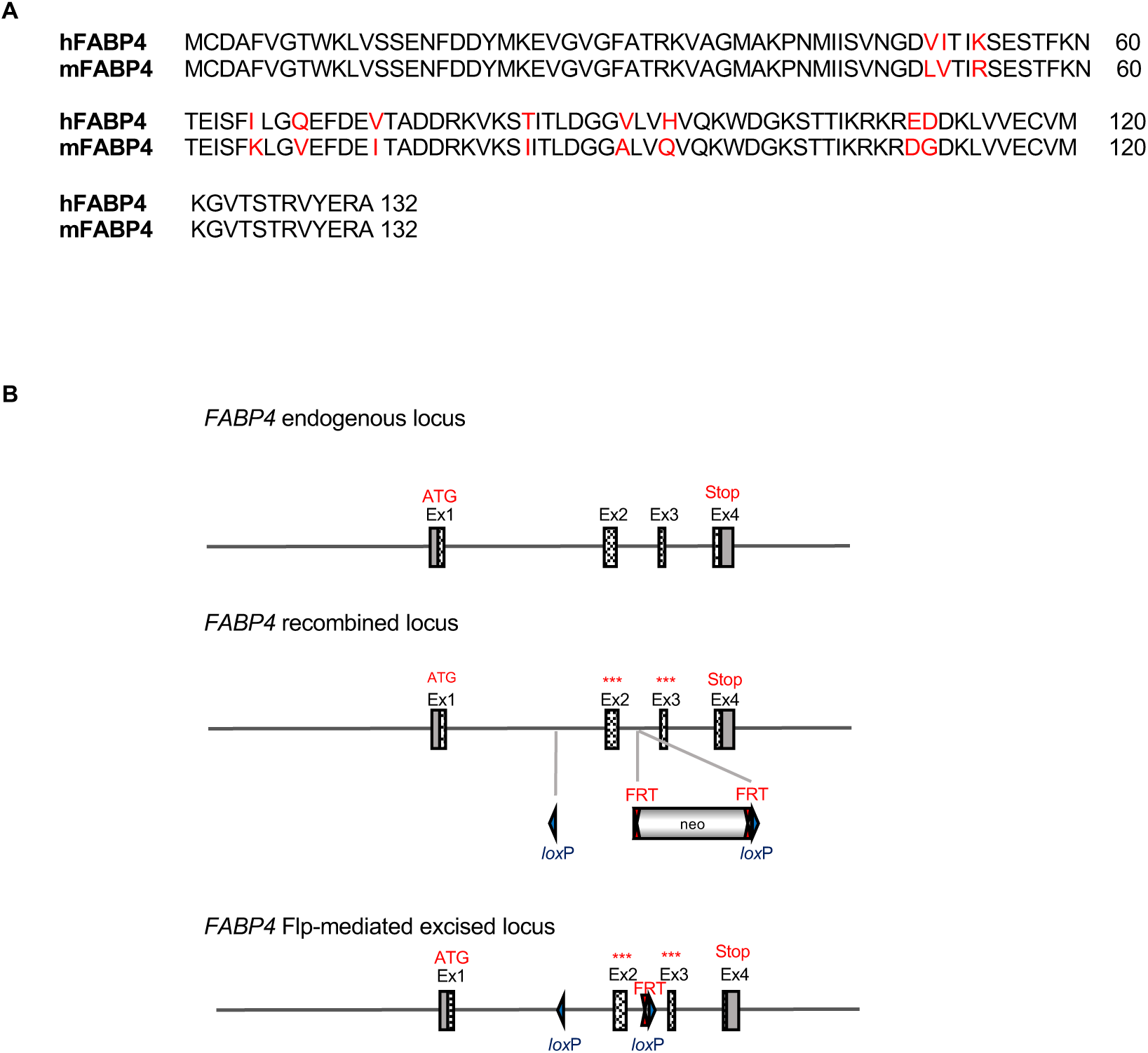
Construct design of humanized FABP4 flox mice. (**A)** The mouse *FABP4* gene was first converted to the the human *FABP4* open reading frame by substituting the 11 amino acids differing between mouse and human *FABP4* located in exons 2 and 3. (**B**) Exon 2 was flanked at the 3’ end by a FRT-neomycin-FRT-loxP cassette and by a single *lox*P site at the 5’ end. This distal *lox*P site was positioned upstream of exon 2 within the intron 1 sequences. The model was generated by homologous recombination in embryonic stem cells. The FRT-flanked selection cassette was removed *in vivo* by crossing with Flp-recombinase-expressing mice. Hatched rectangles represent FABP4 coding sequences. Gray rectangles indicate non-coding exon portions. Solid lines represent chromosome sequences. The 11 murine/human amino acid substitutions are represented by stars.

### Baseline plasma FABP4, FABP5, glucose, and insulin measurements

Baseline plasma measurements were performed in mice ranging from 8-18 weeks of age. Within each replicate experiment, mice were age-matched. Mice were food deprived for 6 hours from 9am to 3pm prior to sample collection. Blood glucose was measured from a tail nick blood sample using a glucometer (Contour Next, Ascensla Diabetes Care). 70µL of blood was collected from a tail nick in non-restrained mice for measurement of plasma FABP4, FABP5, and insulin levels. Due to sensitivity of plasma FABP4 to induction by stress, mice were acclimated to handling in the days prior to blood collection. FABP4 levels obtained from 6h food deprived baseline lipolysis samples were also included in the analysis.

### *In vivo* lipolysis

Lipolysis experiments were performed in mice ranging from 8-18 weeks of age. Within each replicate experiment, mice were age-matched. Mice were food deprived from 9am for 6 hours, and lipolysis was induced by injection of isoproterenol in PBS (10mg/kg, IP, Tocris Bioscience, cat# 1747). 70uL of blood was collected from a tail nick into heparinized capillary tubes at baseline (0) before injection, and 15, 30, and 60 minutes after injection, for measurement of plasma FABP4, insulin, glycerol, and NEFA levels.

To determine the role of circulating FABP4 in the lipolysis-induced insulin response, FABP4-KO mice were administered FABP4 (7µg, IP, recombinant human FABP4, produced in-house) in PBS vehicle or PBS alone 30 minutes prior to induction of lipolysis with isoproterenol. Responses were compared to those of wildtype mice injected with PBS. The FABP4 injection experiment was performed in mice that were wildtype and knock-out for mouse FABP4 (3), on C57BL/6 background.

### Lipolysis in adipose explants

Perigonadal adipose tissue was collected from four 18 week old male mice per group and washed with DMEM containing 10% Cosmic calf serum (CCS-DMEM). They were then transferred to fresh CCS-DMEM and minced into 2-4mm^3^ pieces with scissors. The pieces were washed 5 times with CCS-DMEM and then incubated for 1hr in CCS-DMEM. For each mouse 3 replicates of 8 pieces of adipose tissue were then transferred to 3 wells of a 6 well plate with CCS-DMEM. The plates were incubated for 1h at 37°C, after which conditioned media was collected for measurement of basal secretion of FABP4, glycerol, and NEFA. Next, lipolysis was induced by incubation in 20µM forskolin (FSK) in CCS-DMEM for 1 hour at 37°C, after which the media was discarded. Fresh media with 20µM FSK was added and the explants were again incubated at 37°C for 1h, after which conditioned media was collected for analysis of lipolysis-induced secretion. Secretion was normalized to weight of the explants.

### Adipose SVF and adipocyte isolation

Perigonadal adipose tissue was collected and washed with PBS, and transferred to 10cm plates containing 3mL of ice-cold Krebs-Ringer HEPES, pH 7.4, containing 2% bovine serum albumin (BSA) and 2mg/mL Type 1 collagenase. The tissue was minced to obtain fragments of ∼1mm^3^ and then transferred to 50mL conical tubes and digested at 37°C with shaking at 100rpm. The resulting cell suspension was passed through a 100µm filter and centrifuged at 600g for 10 minutes. The floating adipocytes were transferred to PBS and centrifuged at 600g for 10 minutes. The remaining solution contained the SVFs and was reserved for SVF isolation. After centrifugation the floating adipocytes were removed and transferred to 5mM EDTA in PBS containing 2% BSA and spun at 600g for 1 minute at room temperature. The floating adipocytes were transferred to a 1.5mL tube, centrifuged again at 600g for 1 min at room temperature, and any remaining liquid was removed from the tube. Equal volume of 2X lysis buffer was added to the adipocytes for protein isolation. For SVF isolation, the SVF fraction was washed in PBS and centrifuged at 1700rpm for 10 minutes. The pellet was lysed in 80µL of 1X lysis buffer for protein extraction.

### Primary endothelial cell isolation and culture

Endothelial cells were isolated from liver alone (20 wk old mice) or liver, heart, and lungs (13-14 wk old mice). Mice were deeply anesthetized with 300mg/kg ketamine and 30mg/kg xylazine and transcardially perfused with 25mL of PBS. Tissues were removed and placed in DMEM media with 1% penicillin-streptomycin. Each tissue was then transferred to a 6cm dish with 5mL of digestion buffer consisting of 2mg/mL Type 1 collagenase in 1% BSA in PBS with 1mM CaCl_2_ and 1mM MgCl_2_. Tissues were minced with a blade for 1min and then the tissues from each animal were combined into a 50mL conical tube and placed horizontally in an orbital shaker set at 100rpm, 37°C, for 1 hour. The digested tissues were then strained with a 70µm filter and spun at 1300rpm for 5 minutes at 4°C. The supernatant was removed and the cells were washed twice with 0.1% BSA in PBS, centrifuged at 1300rpm for 5 minutes at 4°C between washes. The resulting pellets were suspended in 1mL of endothelial cell media (Endothelial Cell Growth Kit-VEGF, ATCC cat# PCS-100-030) and counted. 2×10^7^ cells were incubated with 20µL of CD31-coated microbeads (Militenyi Biotec, cat# 130097418) in 180µL of MACS separation buffer (MACS BSA stock solution, cat# 130-091-376; autoMACS Rinsing Solution, cat # 130-091-222, Militenyi Biotec) for isolation of endothelial cells, according to the manufacturer’s instructions. The isolated CD31-positive cells were used for cell culture, collected for FACS analysis, or lysed in RIPA buffer for protein analysis. For determination of FABP4 secretion cells were plated on 6-well 0.1% gelatin-coated plates with endothelial media. For each mouse 3 wells with 300,000 cells/well were plated. The next day the cells were incubated for 4 hours with fresh media, and media and cells were collected for FABP4 measurement and protein quantification. For determination of FABP5 secretion 3 wells per mouse of 30,000 cells/well were plated in 96 well plates and incubated for 24 hours with fresh media before media was collected for FABP5 measurement.

### FACS analysis

For FACS analysis, CD31-positive cells isolated from liver were suspended in 100µL of MACS separation buffer per 10^6 cells. Cells were stained with 2µL of CD31-FITC (mouse, clone: 390) and 2µL of anti-F4/80-APC (mouse, clone: REA126) in the dark at 4°C for 10 min, then washed with 2mL of MACS separation buffer and centrifuged at 300g at 4°C for 10 minutes. Supernatant was removed and the pellet was resuspended in MACS buffer and analyzed by flow-cytometer (BD FACSCalibur Flow Cytometer, BD Biosciences).

### HUVEC cell culture experiments

Primary human umbilical vein endothelial cells (HUVECs, ATCC, cat# PCS-100-010, passage 1) were cultured in growth medium (Vascular Cell Basal Medium, ATCC, cat# PCS-100-030) supplemented with endothelial cell growth kit-VEGF (ATCC, cat # PCS-100-041) in gelatin-coated tissue culture flasks. Passages 3-6 were used for experiments. For experiments, HUVECs were seeded onto 12-well collagen-coated tissue culture plates (Corning 356500) at a density of 1×10^5^ cells/well (day 0) and fed with fresh growth medium every other day. The cells reached 100% confluence around day 3 and “cobblestone” stage around day 7. Intracellular FABP4 protein expression and secretion were measured from days 4 through 14 of culture. FABP4 secretory responses to the following agents were determined on days 11 or 12. Endoplasmic reticulum-Golgi pathway secretion inhibitors: Brefeldin A (Tocris Bioscience, cat# 1231), monensin sodium salt (Tocris Bioscience, cat# 5223). Lysosomal secretion inhibitors: chloroquine (N4-(7Chloro-4-quinolinyl)-N1,N1-dimethyl-1,4-pentanediamine diphosphate salt, Sigma, cat# C6628), ammonium chloride (NH_4_Cl, Boston Bioproducts, cat# P-734). Induction of intracellular calcium: histamine (histamine dihydrochloride, Tocris Bioscience, cat# 3545). Lipolytic stimuli: forskolin (FSK, Cayman Chemical, cat# 11018), 3-isobutyl-1-methylxantine (IBMX, Sigma, cat# 17018), isoproterenol hydrochloride (Tocris Bioscience, cat# 1747), and CL-316,243 disodium salt (Tocris Biosience, cat# 1499). Dose responses to FSK were compared to those of 3T3-L1 differentiated adipocytes, cultured as previously described (14). For treatment, cobblestone HUVECs were washed twice with PBS and incubated in fresh vascular cell basal medium with treatment reagents (dose and duration specified in Figure 7 and supplemental Table 1). At the end of the treatment, the media were collected and spun at 500g for 15 min. The supernatant was used as conditioned medium. The cells were washed twice with cold PBS and lysed for subsequent protein analyses.

### Peritoneal macrophage collection

Mice were injected with thioglycolate (1mL, IP, BD, cat# 221195). Four days later mice were euthanized and 5mL of PBS was injected into the peritoneum. The peritoneal fluid containing macrophages was collected and spun down to pellet the cells. The cell pellet was lysed with 200uL hypertonic lysis buffer (150mM NaCl, 50mM EDTA, 100mM Triton-X-100). The resulting cell lysate was assayed for FABP4 levels by ELISA.

### Islet isolation and insulin secretion

Primary mouse islet isolation and insulin secretion assays were performed as previously described (24). Briefly, 20 islets were hand-picked, washed twice with Krebs Ringer Buffer (KRB) without glucose, and pre-incubated in low glucose (LG; 2.8mM) KRB for 1 hour at 37°C. KRB was then removed and islets underwent successive incubations for 20 minutes at 37°C with 250μL of fresh KRB containing LG, followed by high glucose (HG; 16.7mM), then HG with FSK (10μM), and finally HG with KCl (30mM), to determine maximal insulin secretory capacity. At the end of each incubation supernatant was removed and collected for analysis of insulin levels. Following the stimulation protocol, cells were lysed in 100μL acid ethanol (70% ethanol, 1% HCl in water) and stored at 4°C overnight. Samples were dried with a SpeedVac and resuspended in 60μL ultrapure water. DNA was quantified by Nanodrop (ThermoFisher, USA). Insulin was quantified using insulin HTRF (Insulin Ultrasensitive Assay kit 62IN2PEH; Cisbio, USA) and normalized to DNA content.

### Plasma and media assays

Plasma and media FABP4 were measured by in-house FABP4 ELISA or a commercially available assay (Human Adipocyte FABP (FABP4) ELISA, Biovendor, cat# RD191036200R). For in-house FABP4 ELISA we used anti-FABP4 antibodies produced for the Hotamışlıgil lab by the Dana Farber Antibody Core (clone 351.4.6H7.G3.G9 for capture, HRP-tagged clone 351.4.5E1.H3 for detection) and recombinant human FABP4 as a standard (R&D Systems, cat# DY3150-05). For Figure 8C WT mouse plasma FABP4 was measured using a commercially available assay for mouse FABP4 (Mouse Adipocyte FABP (FABP4) ELISA, Biovendor, cat# RD291036200R). Plasma and media FABP5, insulin, von Willebrand factor (vWF), and endothelin-1 (ET-1) levels were determined by commercially available ELISA (CircuLex Mouse FABP5/-E-FABP/mal1 ELISA kit, cat# CY-8056; Alpco Mouse Ultrasensitive Insulin ELISA, cat# 80-INSMSU-E01 or Crystal Chem Ultra Sensitive Mouse Insulin ELISA, cat# 90080; Human vWF-A2 DuoSet ELISA, R&D Systems, cat # DY2764-05; Human Endothelin-1 QuantiGlo ELISA Kit, R&D Systems, cat # QET00B). Plasma and media NEFA and glycerol were measured by enzymatic colorimetric assays (FujiFilm, Wako HR Series NEFA-HR(2) cat #s 999-34691, 995-34791, 991-34891, 993-35191; Sigma Free Glycerol Reagent, cat# F2468). Media lactate dehydrogenase (LDH) was assayed by enzymatic colorimetric assay (LDH Cytotoxicity Detection Kit, Takara Bio, cat# MK401).

### Protein extraction, SDS PAGE, and Western blotting

Tissues for protein analysis were collected from mice after 6 hour daytime food withdrawal. Western blots for FABP4 and FABP5 were performed in protein lysates from whole tissue or isolated cell populations. To clear tissues of circulating FABP4 and FABP5, mice were intracardially perfused with saline prior to tissue collection. Whole adipose tissues were lysed in in ice-cold RIPA buffer (Cell Signaling Technologies, cat# 9806) containing 2mM activated Na_3_VO_4_ and 1% protease inhibitor cocktail (Sigma, cat# P340). Lysates were assayed for protein concentration by BCA assay (Thermo Fisher/Pierce, cat# 23225), diluted with 5X Laemmli buffer and boiled for 5 minutes at 95°C before being subjected to SDS-polyacrylamide gel electrophoresis using 15% (made in-house) or 4-20% (Criterion TGX Stain-Free Protein Gel, Bio-Rad) gels. Protein was transferred to PVDF membranes which were then blocked for at least 1 hour in TBS-T with 5% protease-free BSA. FABP4 protein was detected using 1:1000 diluted HRP-tagged anti-FABP4 antibody (clone 351.4.5E1.H3, generated for Hotamışlıgil lab by Dana Farber Antibody Core). FABP5 was detected with 1:2000 rabbit anti-FABP5 antibody (Cell Signaling Technology, cat# 39926S) followed by 1:10000 HRP-tagged anti-rabbit secondary antibody (Cell Signaling Technology, cat #7074). HRP signal was detected with chemiluminescent substrate (Pierce SuperSignal West Pico or Femto Plus, Thermo Fisher, cat# 34580 or 34094). Following FABP4 or FABP5 detection, membranes were stripped (Pierce Restore Western Blot Stripping Buffer, Thermo Fisher, cat# 21059) and incubated with HRP-tagged anti-β-tubulin antibody (Abcam, cat# ab21058) to measure β-tubulin as a loading control. Antibodies were diluted in TBS-T with 3% protease-free BSA and incubated overnight at 4°C. Antibodies have been validated with protein lysates from FABP4-KO and FABP5-KO mice as negative controls and recombinant FABP4 and FABP5 as positive controls. The FABP5 was hexahistidine-tagged (Cayman, cat# 10007443), and was 19.3kDa in size instead of the 15kDa expected size for native FABP5.

### Immunohistochemistry

Liver, perigonadal adipose, brown adipose tissue, and pancreas were fixed in 10% zinc formalin overnight at room temperature and then transferred to 70% ethanol. Tissues were embedded in paraffin and sectioned at the Dana Farber Rodent Histopathology Core. Slides were deparaffinized and rehydrated, then heat-induced epitope retrieval was performed by boiling in 10mM citrate buffer, pH 6.0, for 15 minutes, followed by washing with deionized water. For pancreas FABP4 immunostaining endogenous peroxidase activity was blocked by incubating for 5 minutes with 3% hydrogen peroxide in deionized water. Slides were washed, then incubated with serum-free protein block for 5 minutes (Novocastra Protein Block Protein Block Serum Free, Leica Biosysems, cat# RE7102), followed by incubation with rabbit anti-FABP4 polyclonal antibody (adipose, liver: 1:500; pancreas: 1:800; Sigma, cat #HPA002118) in antibody diluent (Agilent, cat#S080983-2) overnight at 4°C. Slides were washed and incubated with anti-rabbit poly-HRP-IgG secondary antibody (Novolink Polymer Detection System, Leica Biosystems, cat# RE7200-CE) for 30 minutes at room temperature. They were then washed again and peroxidase activity was developed with 3,3’-diaminobenzidine (DAB) chromogen and peroxidase substrate, and counterstained with hematoxylin.

For pancreas islet insulin immunostaining, 3 serial sections per pancreas at 250μm apart were processed. Immunostaining was performed as described previously (24). Images of each section were acquired using Aperio Imagescope at 20x magnification. The beta cell area was calculated by using positive pixel count analysis (Aperio ImageScope). Islet number and size were determined by manually circling insulin positive clusters in the Aperio ImageScope software, and was performed blinded to the sample IDs.

### Statistics

Statistical analysis was performed using GraphPad Prism version 9.4.0 for MacOS, GraphPad Software, San Diego, California USA, www.graphpad.com. Statistical methods are indicated in the figure legends. All data are presented as mean ± SEM.

### Study approval

All *in vivo* mouse studies were approved by the Harvard Medical Area Standing Committee on Animals.

## Author contributions

K.E.I designed and performed the *in vivo* experiments, analyzed the data, and prepared the manuscript. A.L., K.J.P., C.D-G, M.C. and G.Y.L. assisted with *in vivo* experiments. K.J.P. and A.L. designed and performed the primary endothelial experiments. K.J.P. designed and performed *in vitro* islet experiments. G.Y.L. designed and performed the HUVEC experiments. A.L., C.D-G., and M.C. performed tissue and plasma assays. K.J.P., A.L., and G.Y.L. analyzed the data and revised the manuscript. G.S.H. conceived, supervised and supported the project, designed experiments, interpreted results, and revised the manuscript.

## Acknowledgments

The authors thank members of the Hotamışlıgil laboratory, past and present, for their contributions to our understanding of FABP4 and their helpful discussions, and special thanks to those who assisted with experiments. FABP4 floxed mice used in this study were developed by GenOway. This research is supported by the Sabri Ülker Center for Metabolic Research, the Juvenile Diabetes Research Foundation (JDRF; 2-SRA-2016-147-Q-R, 2-SRA-2019-660-S-B), and National Institutes of Health (RO1 DK123458). A JDRF Postdoctoral Fellowship (3-PDF-2017-400-A-N) funded K.J.P.

## Notes

### Competing Interest Statement

G.S.H. is in the Scientific Advisory Board of Crescenta Biosciences and holds equity. The Hotamışlıgil lab holds intellectual property related to hormonal FABP4 and its therapeutic targeting. Other authors have no conflicts of interest to declare.

