## Supplementary Figures for "Endothelial FABP4 constitutes the majority of basal circulating hormone levels and regulates lipolysis-driven insulin secretion"

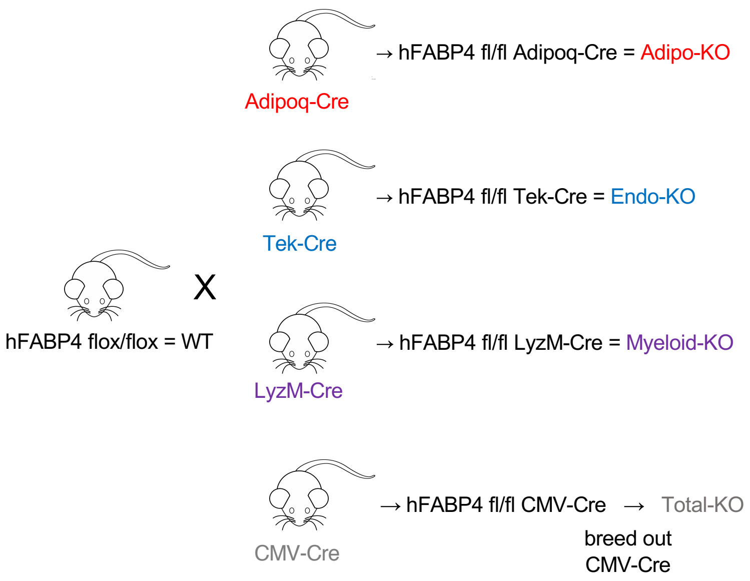


**Supplementary Figure 1: Breeding scheme for tissue-specific FABP4 deletion mice.** Mice with floxed humanized FABP4 (hFABP4 flox/flox) were crossed with Adiponectin-Cre, Tek-Cre, LyzM-Cre, or CMV-Cre mice to generate mice with deletion of FABP4 in adipocytes (Adipo-KO), endothelial cells (Endo-KO), myeloid cells (Myeloid-KO), or the whole body (Total-KO). Since crossing with CMV-Cre produces germline deletion of FABP4, CMV-Cre was bred out of the Total-KO mouse line. hFABP4 flox mice are referred to as WT. WT mice used in this study are littermates of Adipo and Endo-KO mice that did not express Cre recombinase.

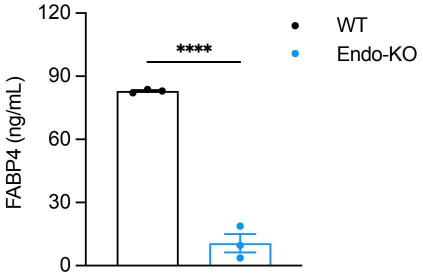


**Supplementary Figure 2**: Peritoneal macrophage lysate FABP4 levels from WT and Endo-KO mice. ****p<0.0001. Data were analyzed by unpaired t-test and are presented as mean ± SEM.

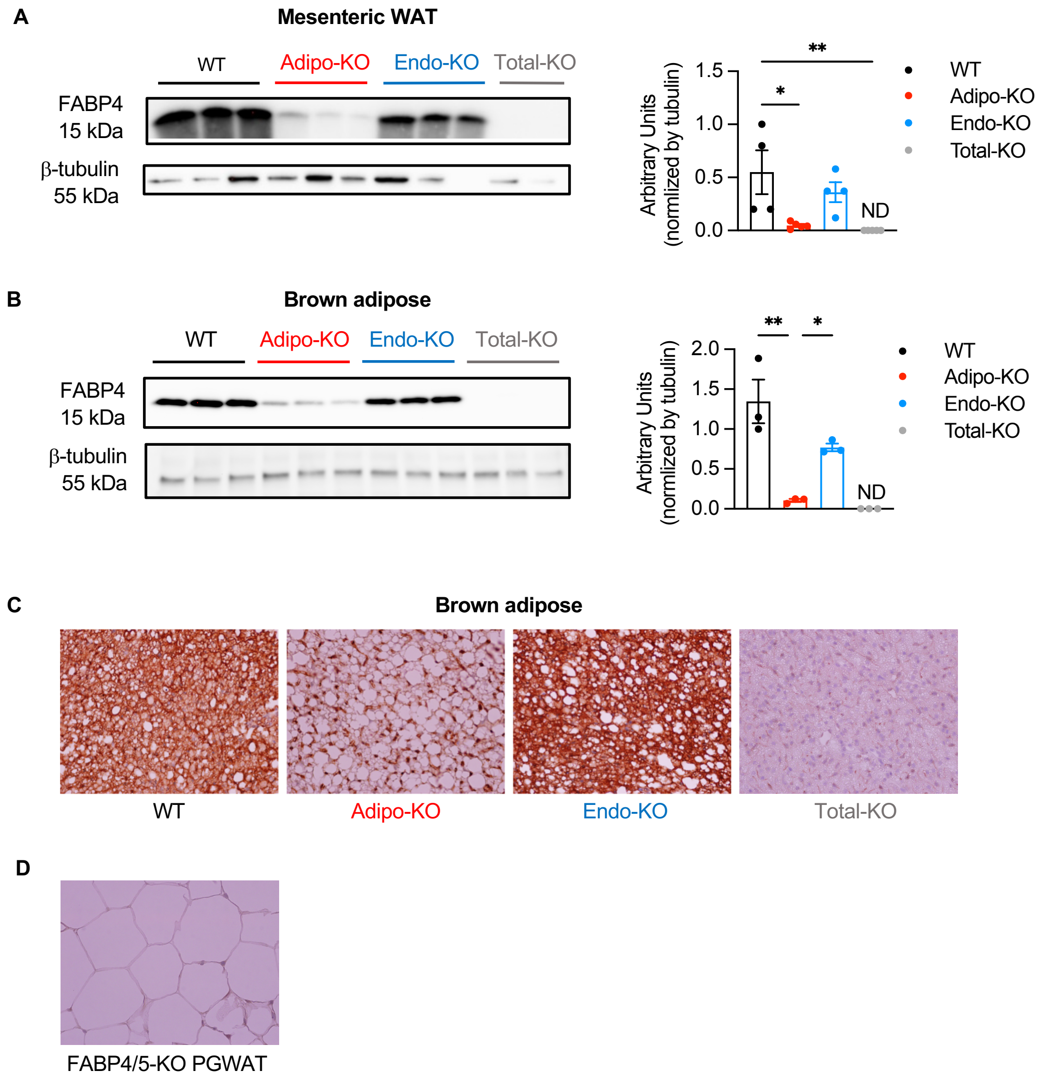


**Supplementary Figure 3:** (**A**) Representative immunoblot of FABP4 expression vs. β-tubulin loading control and quantification of FABP4 relative to β-tubulin signal in mesenteric adipose tissue of WT, Adipo-KO, Endo-KO, and Total-KO mice. **p<0.005. *p<0.05. (**B**) Immunoblot of FABP4 expression vs. β-tubulin loading control and quantification of FABP4 relative to β-tubulin signal in brown adipose tissue of WT, Adipo-KO, Endo-KO, and Total-KO mice. (**C**) FABP4 immunostaining in brown adipose tissue from WT, Adipo-KO, Endo-KO, and Total-KO mice. 40X magnification. (**D**) FABP4 immunostaining in perigonadal adipose tissue (PGWAT) from FABP4/5-KO mice. 40X magnification. Data were analyzed by one-way ANOVA followed by Tukey's multiple comparison test and are presented as mean ± SEM. ND: No signal detected.

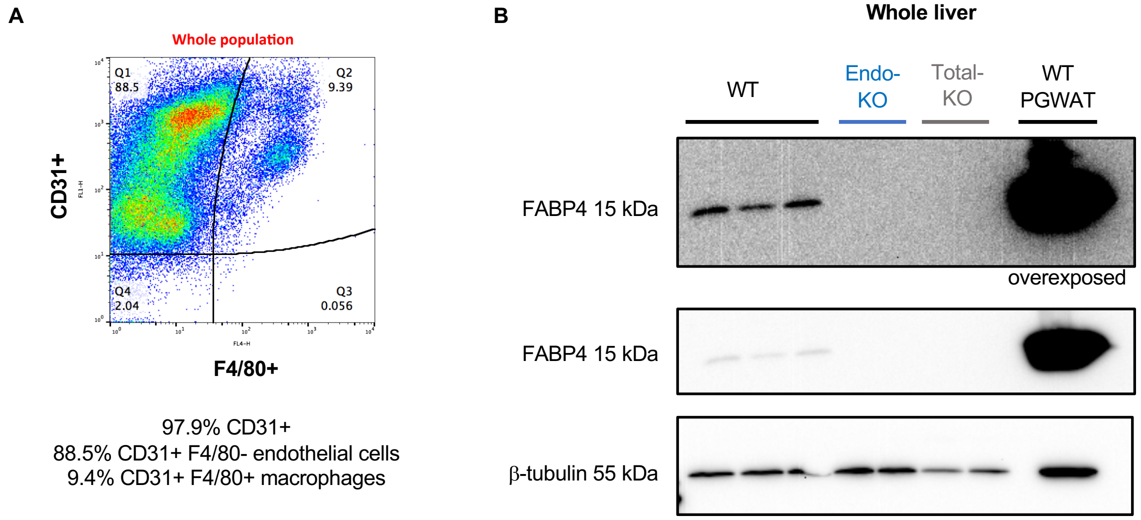


**Supplementary Figure 4:** (**A**) FACS analysis of WT liver endothelial cells isolated with anti-CD31antibody coated beads, and co-stained with anti-CD31-FITC and anti-F4/80-APC to determine the percentage of F4/80-positive macrophages within the CD31 population. (**B**) Immunoblot of FABP4 expression in liver lysates from WT, Endo-KO, and Total-KO mice vs. β-tubulin loading control. Lysates from PGWAT of WT mice included as a positive control.

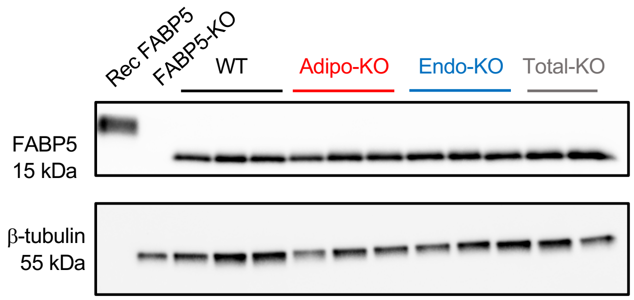


**Supplementary Figure 5.** Immunoblot of perirenal adipose FABP5 expression in WT, Adipo-KO, Endo-KO, and Total-KO mice vs. β-tubulin loading control. Recombinant FABP5 was used as a positive control and FABP5 KO adipose lysate as a negative control. Recombinant FABP5 is hexahistidine-tagged and travels at 19kDa, compared to endogenous FABP5 at 15 kDa.

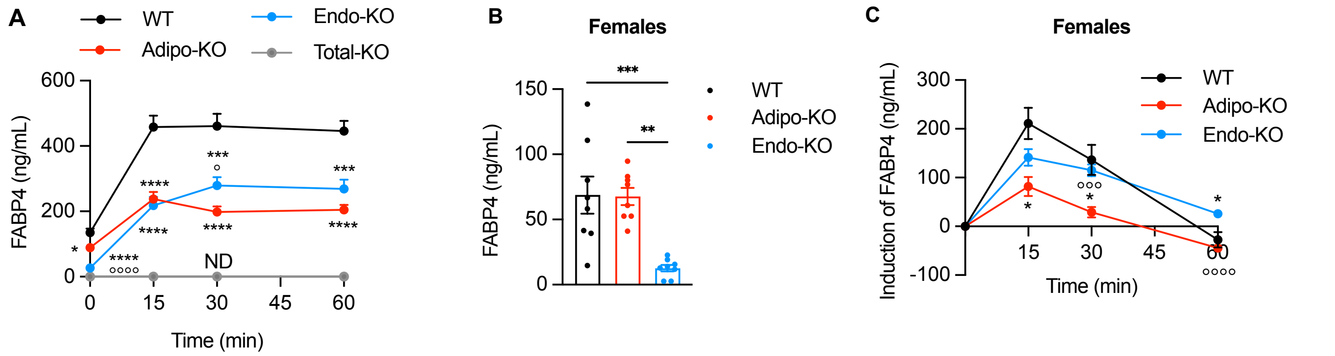


**Supplementary Figure 6.** (**A**) Plasma FABP4 responses to 10mg/kg isoproterenol-induced lipolysis in male WT, Adipo-KO, Endo-KO, and Total-KO mice. Data are pooled from 6 experiments. WT n=52, Adipo-KO n=39, Endo-KO n=34, Total-KO n=8. ****p<0.0001, ***p<0.001, *p<0.05 vs. WT.°°°°p<0.0001, °p<0.05 vs. Adipo-KO. (**B**) Baseline plasma FABP4 levels from S5C lipolysis experiment in female WT, Adipo-KO, Endo-KO, and Total-KO mice. **p<0.005, **p<0.001 vs. WT. (**C**) Plasma FABP4 responses over baseline to 10mg/kg isoproterenol-induced lipolysis in female WT, Adipo-KO, and Endo-KO mice. n=8/group. *p<0.05 vs. WT. °°°°p<0.0001, °°°p<0.001 vs. Endo-KO. S6A and S6C were analyzed by repeated measures 2-way ANOVA with Geisser-Greenhouse correction, followed by Tukey's multiple comparison test. S6B was analyzed by one-way ANOVA followed by Tukey's multiple comparison test. Data are presented as mean ± SEM. ND: No signal detected.

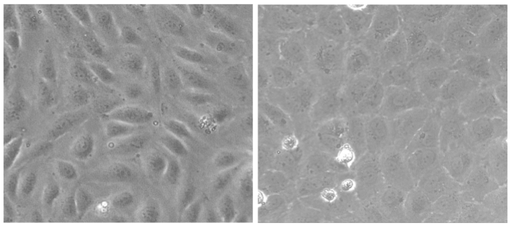


**Supplementary Figure 7.** Light microscopy image of HUVECs at days 4 and 11 post-seeding. Day 11 HUVECs demonstrate “cobblestone” phenotype. 10X magnification.

| | | Reagent |
|---|---|---|
| **Secretory Mechanisms** | **ER-Golgi pathway inhibition** | Brefeldin A, monensin (Up to 20uM) |
| | **Lysosomal pathway inhibition** | Chloroquine (Up to 50uM) $NH_4Cl$ (Up to 50mM) |
| | **Increased intracellular Ca++** | Histamine (Up to 25uM) |
| **Stimuli** | **Lipolytic/Adrenergic** | FSK (Up to 25uM), IBMX (1mM), CL-316,243 (Up to 20uM), isoproterenol (Up to 50uM) |

**Supplementary Table 1: Agents that did not affect HUVEC FABP4 secretion**. Day 12 HUVECs were treated for 2 hours with agents known to target secretory mechanisms in adipocytes and endothelial cells and stimulate adipocyte FABP4 secretion.
